# Single-Cell Transcriptomic Analysis of mIHC Images via Antigen Mapping

**DOI:** 10.1101/672501

**Authors:** Kiya W. Govek, Emma C. Troisi, Zhen Miao, Steven Woodhouse, Pablo G. Camara

## Abstract

Highly-multiplexed immunohistochemistry (mIHC) enables the staining and quantification of dozens of antigens in a tissue section with single-cell resolution. However, annotating cell populations that differ little in the profiled antigens or for which the antibody panel does not include specific markers is challenging. To overcome this obstacle, we have developed an approach for enriching mIHC images with single-cell RNA-seq data, building upon recent experimental procedures for augmenting single-cell transcriptomes with concurrent antigen measurements. Spatially-resolved Transcriptomics via Epitope Anchoring (STvEA) performs transcriptome-guided annotation of highly-multiplexed cytometry datasets. It increases the level of detail in histological analyses by enabling annotation of subtle cell populations, spatial patterns of transcription, and interactions between cell types. More generally, it enables the systematic annotation of cell populations in cytometry data. We demonstrate the utility of STvEA by uncovering the architecture of poorly characterized cell types in the murine spleen using published highly-multiplexed cytometry and mIHC data.

## Introduction

Recently developed technologies for digital imaging and mIHC^1-6^ are enabling the field of histology to enter into a quantitative era, allowing for more complex descriptions of tissue architecture. Imaging mass cytometry^5^, multiplexed ion beam imaging^6^, and co-detection by indexing^2^ (CODEX) can be used to simultaneously profile the expression level of dozens of proteins in a tissue section with single-cell resolution. Despite this progress, the amount of cell types and states that can be simultaneously identified by mIHC is limited. Computational methods for automated identification of cell populations cluster cells according to expression similarities of the profiled antigens. These clusters are then manually annotated using previous knowledge of cell markers. However, this process is generally partial, subjective, and biased^7^. The groups of cells that result from clustering algorithms often differ little in their antigenic profile and the interpretation of those differences is unclear. Moreover, the design of comprehensive antibody panels that include specific markers for every cell type and state present in the tissue is usually unfeasible. Consequently, the amount of annotated cell populations in mIHC analyses is often substantially smaller than the number of clusters produced by automated methods.

To overcome these limitations and improve the annotation of mIHC data, we propose an approach for enriching mIHC slides with single-cell RNA sequencing (RNA-seq) data. Currently available single-cell RNA-seq technologies can profile the expression level of thousands of genes in each cell, allowing for fine classification of cells based on their gene expression profile. Some of the most recent approaches, like CITE-seq^8^, REAP-seq^9^, and Ab-seq^10^, allow for augmenting single-cell transcriptomes with concurrent protein measurements by staining single-cell suspensions with oligo-tagged antibodies. These approaches can therefore be used to determine the quantitative relation between gene and antigen expression levels. Here, we build upon CITE-seq and computational methods for the integration of single-cell omics data^11, 12^ to identify and annotate cell populations in mIHC images (or more broadly, in highly-multiplexed cytometry datasets) based on single-cell gene expression data of the same tissue. Our method, STvEA, consists of three major steps. First, it computationally consolidates the protein expression spaces of the mIHC dataset and a matching CITE-seq dataset using a shared antibody panel. This consolidated protein expression space is used to transfer features (e.g. mRNA cell type assignments, gene expression profiles, etc.) from the CITE-seq dataset into the mIHC images. STvEA then finds an optimal clustering of the CITE-seq mRNA expression data such that the resulting cell populations can be accurately mapped into the mIHC images based on their antigenic profile. In this way, STvEA enables the identification and annotation of cell populations in the mIHC data and the study of spatial patterns of transcription.

We use the murine spleen as a test system to benchmark the stability and performance of STvEA, since well-established antibody panels and high-quality mIHC datasets are readily available for this organ. For that purpose, we have generated a high-quality CITE-seq atlas of the murine spleen and used it with STvEA to annotate published mIHC and mass cytometry datasets of this organ. Our results reveal that STvEA substantially increases the level of phenotypic annotation of these datasets, and enables new analyses of highly-multiplexed cytometry data. In addition, by systematizing the annotation of cell populations, it improves the reproducibility of the results. We have made this approach available to the entire community as open source software (Online Methods).

## Results

### A high-quality CITE-seq atlas of the murine spleen

We aimed to improve and automate the annotation of a published high-resolution mIHC dataset of the murine spleen recently generated with the CODEX technology^2^. CODEX employs an in-situ polymerization indexing procedure to measure the spatial distribution of a panel of protein markers with sub-micrometer resolution. To be able to more accurately annotate this dataset, we generated a high-quality CITE-seq dataset of the murine spleen using the same 30-antibody panel (Supplementary Table 1) and mice matching those of the CODEX dataset. In total, we profiled the transcriptome and antigen levels of 7,097 cells (with ≥ 1,200 mRNA UMIs) using CITE-seq. The median Spearman correlation among the observed expression of mRNAs and the proteins they code for was 0.32, consistent with previous CITE-seq studies^8^. We used single-cell variational inference (scVI)^13^ to obtain a latent space representation of the mRNA data and clustered the cells in this space using an in-house consensus algorithm (Online Methods). Our analysis found 17 clusters and no noticeable batch effects (Figs. 1a, Supplementary Fig. 1). We performed differential expression analysis to annotate the clusters based on the expression of known marker genes (Fig. 1b, Supplementary Table 2). Additionally, we utilized a spectral graph method^14, 15^ to characterize the transcriptomic heterogeneity that originates from the continuous maturation processes occurring in the spleen (Fig. 1c, Supplementary Table 3). This approach allowed us to identify genes with significant gradients of expression within one or several clusters, and we used these results to further annotate the atlas. Overall, we identified 30 cell populations, comprising most of the known splenic cell types^16, 17^ (Fig. 1a). These results represent a substantial increase in resolution with respect to previous single-cell RNA-seq atlases of the murine spleen^18-20^.

**Figure 1.**
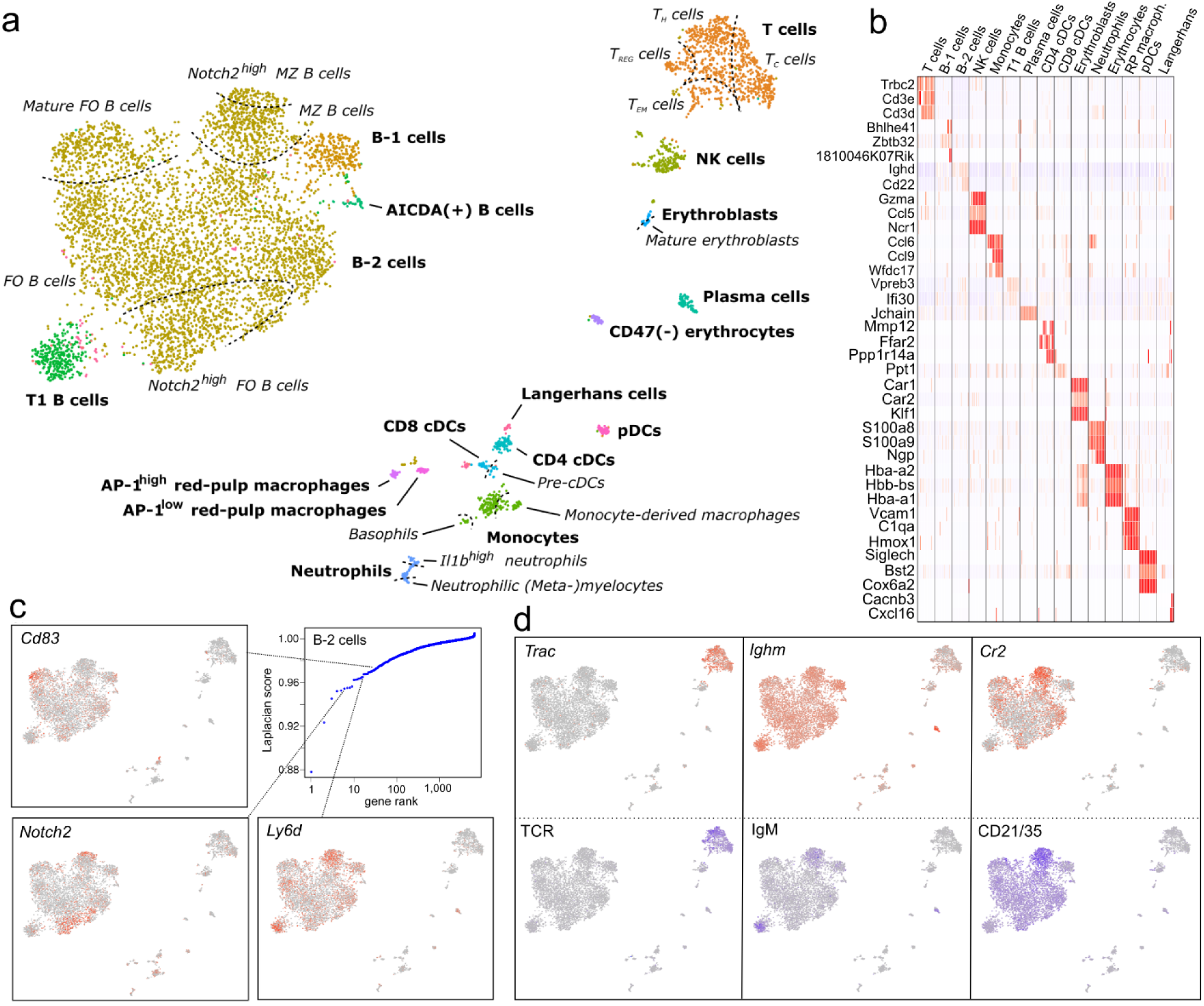
A high-resolution CITE-seq atlas of the murine spleen. **a)** UMAP representation of the mRNA expression data of 7,097 cells from the murine spleen profiled with CITE-seq. Cell populations were identified by clustering (represented in different colors) and annotated by differential expression analysis (bold text) and a spectral graph method (italic text; see also panel c). Dashed lines represent soft transitions in the transcriptome of cells. **b)** Heatmap depicting the expression of some of the top differentially expressed genes in each cluster. **c**) Analysis of the cellular heterogeneity within the clusters of B-2 cells using a spectral graph approach. Genes were ranked according to their Laplacian score and the statistical significance was assessed for each gene by randomization. In the figure, the expression levels of some of the significant genes are depicted in the UMAP representation. The complete results are provided for all clusters in Supplementary Table 3. **d)** mRNA expression levels of *Cr2, Ighm*, and *Trac* (top) and the expression levels of the proteins they code for (bottom).

### Mapping of the splenic CITE-seq atlas onto histology sections profiled with CODEX

We noticed that most of the cell populations identified in the transcriptomic analysis of the CITE-seq atlas were also localized in the protein expression space (Supplementary Fig. 2). This observation indicates that small differences in cellular epitope levels are often representative of distinct cell populations, even if those differences do not lead to discrete clusters in the protein expression space. Consequently, we reasoned that mapping the CODEX protein expression space into the CITE-seq protein expression space would allow us to survey the CODEX images for the cell populations identified in the transcriptomic analysis. To lessen the technical differences and facilitate the integration of the two spaces, we devised a common approach to background removal and normalization for CODEX and CITE-seq protein expression measurements (Supplementary Fig. 3). In each dataset, we modeled the distribution of protein levels using a two-component mixture model (Online Methods). Our approach led to improved and more consistent protein expression levels across the two datasets (Supplementary Fig. 3). We then employed a mutual nearest neighbors anchoring strategy^*11, 12*^ to consolidate the signal component of the two datasets into a common protein expression space (Fig. 2a, Online Methods). By looking at the CODEX neighbors of each CITE-seq cell in the consolidated protein expression space, we were able to identify groups of cells in the mIHC images with similar antigenic profiles to those in the CITE-seq dataset, substantially extending the phenotypic annotation of cell types in the CODEX data. Overall, STvEA led to the annotation of 73% (*n* = 57,819) of the cells present in the CODEX mIHC images. It correctly recapitulated the known spatial distribution of splenic cell populations, including the partitioning between red-pulp, B cell zones, and T cell zones, the location of plasmacytoid dendritic cells (pDCs) in T cell zones, the location of monocytes in the red-pulp, and the positioning of CD4 conventional dendritic cells (cDCs) along the bridging channels that connect T cell zones and the red pulp^21^ (Fig. 2b). Cell populations that were not annotated by STvEA mostly consisted of stromal cells (CD31+ or ERTR7+ cells) with no representation in the CITE-seq atlas due to the non-enzymatic procedure we employed for tissue dissociation. CODEX cell population assignments were consistent across CITE-seq replicates (Fig. 2c, median Pearson’s correlation *r* = 0.998, *p*-value < 10^−10^), and the inferred relative spatial distributions were reproducible across multiple spleens profiled with CODEX (Supplementary Fig. 4).

**Figure 2.**
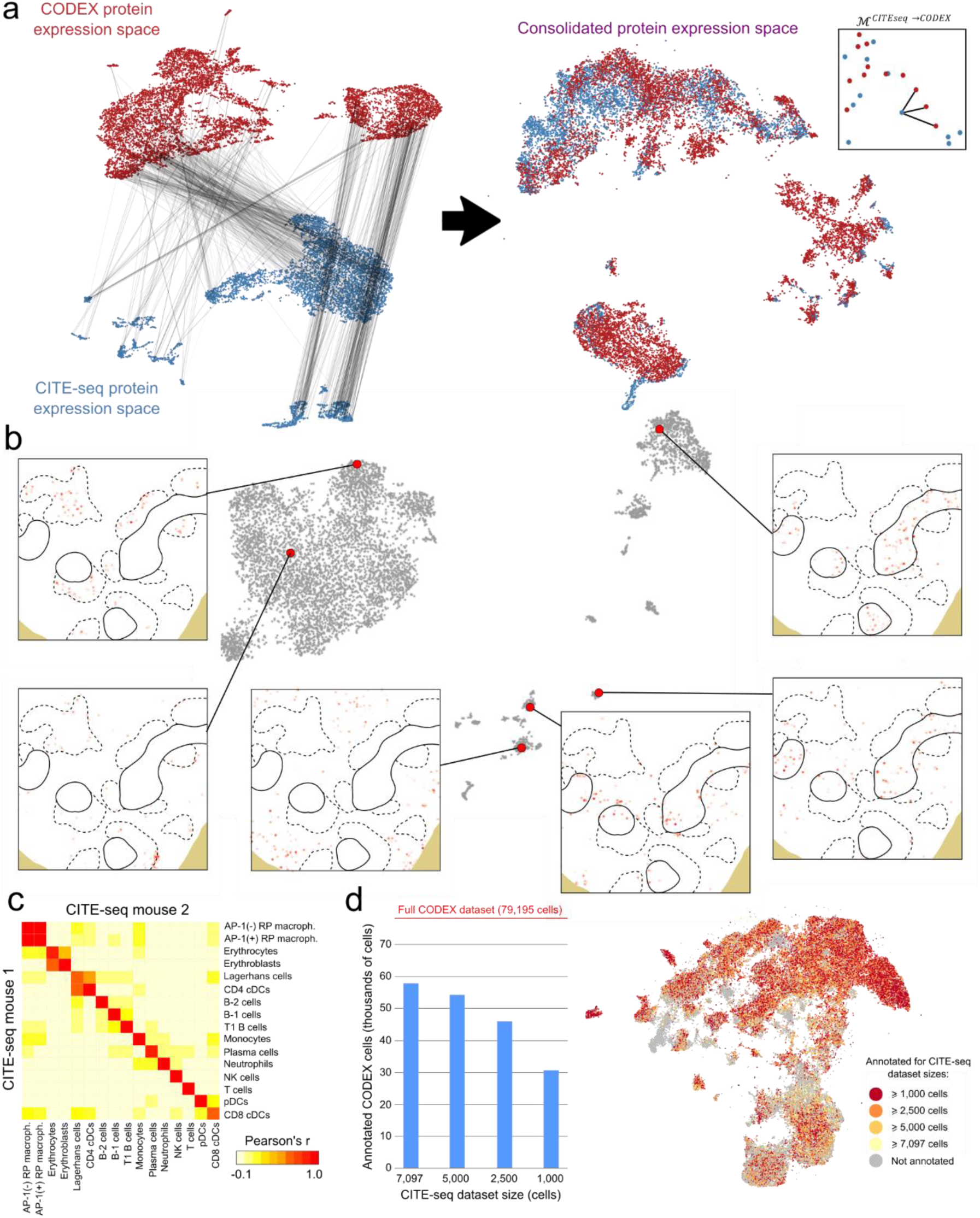
Mapping of the splenic CITE-seq atlas into histology sections profiled with CODEX. **a)** Schematics of the procedure to map the CODEX and CITE-seq protein expression spaces. A set of anchors are identified using a mutual nearest neighbors approach and weighted according to their degree of consistency with the mRNA expression space (left). The anchors are used to merge the CODEX and CITE-seq protein expression spaces into a common space (middle). The transfer matrix ℳ^*CITEseq*→*CODEX*^ is built by looking at the nearest CODEX cells to each CITE-seq cell in the merged protein expression space (right). **b)** Mapping of cells from the splenic CITE-seq atlas into a murine splenic section profiled with CODEX. The figure shows the locations of cells in the splenic section with antigenic profiles similar to those of 6 cells from the CITE-seq atlas. For reference, the T and B cell zones in the tissue section are indicated with solid and dashed lines, respectively. As can be seen in the figure, transcriptomic differences between cells of the same cell type often correspond to different spatial locations. These differences were consistent when the same cells were mapped to other murine splenic sections profiled with CODEX (Supplementary Fig. 3). **c)** Consistency between the annotations of two different murine spleens profiled by CITE-seq into the same CODEX dataset. A heatmap showing the correlation between the CODEX cell assignments for each cell population in the two mice profiled with CITE-seq. In an ideal scenario, diagonal entries would be perfectly correlated (Pearson’s *r* = 1) and off-diagonal entries anti-correlated (Pearson’s *r* < 0). Departures from that situation quantify mapping inaccuracies. As represented in the figure, STvEA has an excellent performance for most splenic cell populations, with the most notable inaccuracies being between AP-1^high^ and AP-1^low^ red-pulp macrophages, and erythrocytes and erythroblasts. **d)** Number of annotated cells in the CODEX dataset as a function of the size (number of cells) of the CITE-seq atlas. The cells annotated by STvEA are indicated in a UMAP representation of the CODEX protein expression space for different sizes of the CITE-seq atlas.

### Quantification of mapping uncertainties and stability

To quantitatively assess the magnitude of mapping errors, we looked at the protein expression profile of cells in the CODEX dataset related to the same CITE-seq cells. The average Pearson’s correlation coefficient between the protein expression profiles of CODEX cells related to the same CITE-seq cell was 0.74. This value varied substantially across the CITE-seq atlas (Supplementary Fig. 5), with cells annotated as B-1 B cells, T1 B cells, and dendritic cells having the lowest correlation coefficients (average Pearson’s *r* = 0.59, 0.55, and 0.56, respectively). However, these coefficients were substantially larger than the correlation between the protein expression profiles of randomly chosen CODEX cells (Supplementary Fig. 5, mean Pearson’s correlation coefficient *r* = 0.25).

We next evaluated the stability of STvEA with respect to the number of cells in the CITE-seq atlas. We randomly sampled cells from the CITE-seq dataset to generate smaller datasets and applied STvEA independently to each of these atlases. The percentage of CODEX cells annotated by STvEA decreased from 73% when using the entire CITE-seq atlas (7,097 cells) to 38% when using only 1,000 cells (Fig. 2d). Expectedly, regions of the CODEX protein expression space with low mapping scores (Online Methods) or less representation in the CITE-seq data were more sensitive to the size of the atlas (Supplementary Fig. 6). However, annotations were highly consistent across atlases of different sizes (median Pearson’s correlation coefficient between the predicted gene expression profile of a CODEX cell when using the original CITE-seq dataset or a down-sampled version with 1,000 cells *r* = 0.93). These results thus indicate that the size of the CITE-seq atlas mainly affects the percentage of annotated cells in the mIHC images, but not the quality of the annotations.

Finally, we assessed the stability of the annotations against changes in the size of the antibody panel. We performed logistic lasso regression using the CITE-seq cell populations as response variable to order the antibodies from least to most informative (Supplementary Fig. 7a, Online Methods). We then successively reduced the size of the antibody panel and applied STvEA. The annotations in the initial analysis were relatively stable against reducing the size of the antibody panel (Supplementary Fig. 7b). In particular, 73% of the annotated cells in the original analysis were still annotated when using a panel with only 9 antibodies, and the median Pearson’s correlation coefficient between cell assignments in these two analyses was 0.99, indicating a large degree of consistency for the annotations. Moreover, all the cell types identified in the mRNA analysis were still well-represented in the annotations of the CODEX dataset when using 9 antibodies (Supplementary Fig. 7c). Based on these results we conclude STvEA can provide a high level of phenotypic annotation with relatively small antibody panels, as long as antibodies are suitably chosen and cell populations are represented in the reference CITE-seq atlas.

### Optimized annotation of cell populations

The analysis described above shows that mapping accuracy is not uniform across the CITE-seq atlas (Supplementary Fig. 5). We therefore reasoned that defining cell populations based exclusively on clustering the mRNA data, without taking into account mapping accuracy, could lead to suboptimal annotation of the mIHC images. In particular, some of the mRNA clusters might not be accurately mapped based on their protein expression profile, whereas other clusters might be split into smaller pieces that could be still accurately distinguished based on their protein expression profile. To overcome this limitation and define cell populations that can be optimally mapped into the mIHC data, we devised a clustering approach of the single-cell mRNA data that takes into account mapping accuracy (Online Methods). Specifically, we use the algorithm HDBSCAN^22^ to establish a simplified hierarchical tree of cell populations based on the single-cell mRNA data. A non-uniform cut of this tree is chosen based on the modularity of the resulting populations in the CODEX protein expression space upon STvEA mapping (Online Methods). This approach partitioned the CITE-seq atlas into 17 phenotypically distinct cell populations that could be accurately mapped onto the mIHC images (Fig. 3a-c). These automated annotations were consistent with those resulting from manual annotation of the CODEX data^2^ (Supplementary Fig. 8), and included several populations that were not identified in the manual analysis, such as pDCs, different stages of erythrocyte maturation, and several B cell subpopulations.

**Figure 3.**
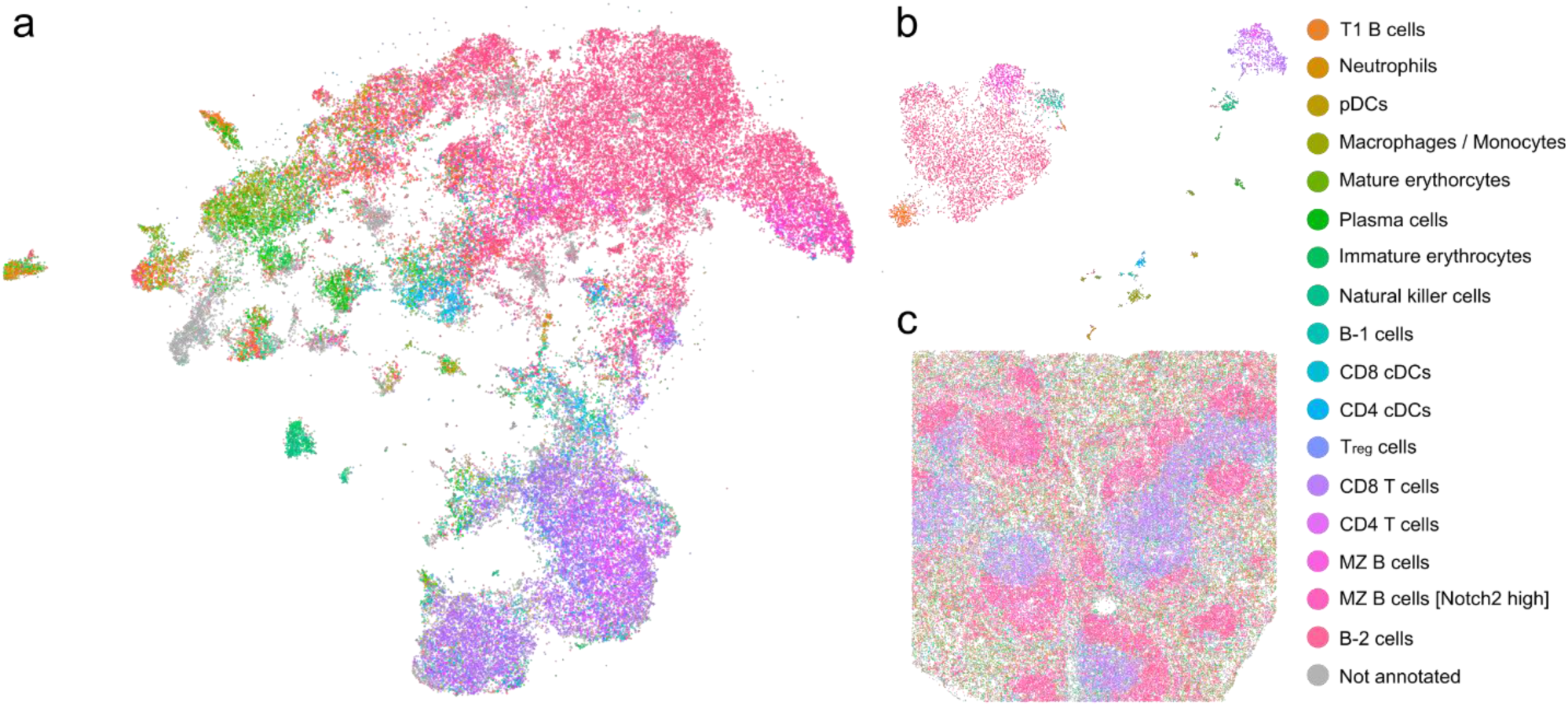
Transcriptome-guided annotation of cell populations in histology sections profiled with mIHC. **a)** UMAP representation of the protein expression space a splenic tissue section profiled with CODEX. The representation is labelled by the cell populations identified by STvEA. In total, 17 phenotypically distinct populations were determined in the mRNA CITE-seq data based on their gene expression profile and their mapping into the CODEX dataset. **b, c)** UMAP representation of the CITE-seq gene expression space (b) and image of the tissue section (d) labeled by the 17 cell populations annotated by STvEA.

We also explored the utility of STvEA as a tool for annotating the clusters produced by some of the existing algorithms for cytometry data analysis, including X-shift^23^, SPADE^24^, and PhenoGraph^25^ (Supplementary Fig. 9). The cell clusters produced by these methods were generally composed of one or two predominant cell types according to the annotations of STvEA. Taken together, these results show that STvEA is a useful and robust tool for the annotation of mIHC data.

### Prediction of spatially-resolved gene expression patterns

The mapping of single-cell transcriptomic data onto mIHC images provided by STvEA allowed us to investigate the predicted spatial patterning of any gene in the mRNA dataset (Fig. 4a). To validate some of the spatially-resolved gene expression profiles predicted by STvEA, we performed multiplexed RNA fluorescent in situ hybridization^26^ (FISH) of several marker genes identified in the differential expression analysis (Fig. 4a, Supplementary Figs. 10 and 11). Specifically, we carried out hybridizations for *Bhlhe41*, a transcriptional repressor highly expressed by B-1 cells^27^ as they mature and migrate from B cell zones into the red pulp^28^; and *Il1b*, expressed by several subpopulations of cDCs, monocytes, macrophages, and neutrophils in the red pulp and T cell zones, but not expressed in B cell zones. In both cases, FISH correctly recapitulated the expression patterns predicted by STvEA (Fig. 4a, Supplementary Figs. 10 and 11), confirming the utility of our computational approach to label mIHC images by gene expression levels. Using this approach, we were also able to resolve in the mIHC images some of the cell differentiation processes that take place in the spleen. For example, the predicted expression patterns of *Car1*, a marker of immature erythroblasts^29, 30^, and *Gypa*, a marker of intermediate and late stages of erythroblast maturation^30, 31^, represented the maturation of erythroblasts in erythroblastic islands of the red pulp^32^ and allowed us to annotate erythroblasts in the CODEX dataset according to their stage of maturation (Supplementary Fig. 12).

**Figure 4.**
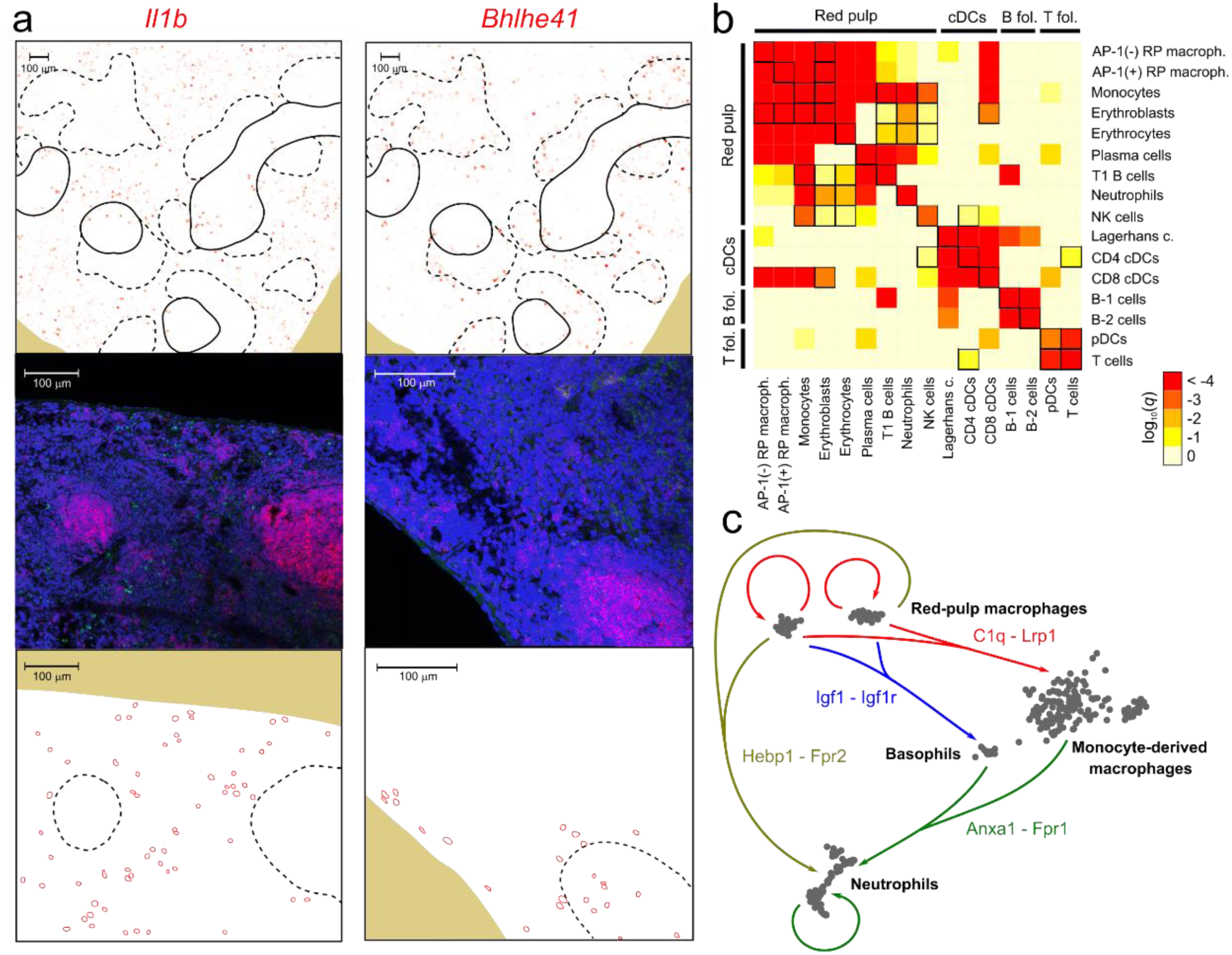
Identification of spatially-resolved gene expression patterns and interactions between cell populations. **a)** mRNA expression levels predicted by STvEA (top) and measured by RNA FISH (middle and bottom) in murine splenic sections for the genes *Il1b* (left) and *Bhlhe41* (right). Red: *Cd79a*, green: *Il1b* / *Bhlhe41*, blue: DAPI. T and B cell zones in the tissue sections are indicated with solid and dashed lines, respectively. We used B cell zones, highlighted by the expression of *Cd79a*, as a reference for comparisons between RNA FISH and CODEX tissue sections. The relative location of cells expressing *Il1b* and *Bhlhe41* with respect to B cell zones is indicated at the bottom. **b)** Identification of interactions between splenic cell populations. Heatmap showing the significance of the spatial co-localization of splenic cell populations, inferred by STvEA. Significant relations (*q*-value ≤ 0.05) that cannot be explained by mapping errors (95% CL) are indicated with black squares. **c)** Some of the significant potential paracrine interactions among red-pulp macrophages, basophils, neutrophils, and monocyte-derived macrophages in the red pulp. Interactions were inferred based on the differential expression of the genes encoding for the ligand and receptor and on their spatial co-localization.

### Identification of cell population interactions

Characterizing interactions between cell populations within the context of tissues is a key step towards understanding cell function. Inferring the cell type of individual cells in mIHC images enabled us to survey candidate interactions between cell populations, further expanding the scope of conventional mIHC image analyses. We devised a graph-based approach for assessing the spatial co-localization of cell populations identified in the transcriptomic analysis while accounting for mapping uncertainties (Online Methods). Significant co-localization patterns recapitulated the well-established immune cellular architecture of the spleen, partitioned into red pulp, B cell zones, and T cell zones (Fig. 4b). T cells, pDCs, and CD4 cDCs were recurrently in close proximity within T cell zones. Similarly, red pulp macrophages, erythrocytes, neutrophils, and monocytes were recurrently in close proximity within the red pulp. In addition, several cell populations showed co-localization patterns that spanned multiple splenic compartments (Fig. 4b). Specifically, CD4 cDCs appeared recurrently in close proximity with T cells in T cell zones and with NK cells in the red pulp (Fig. 4b). These inferred relations were reproducible across multiple spleens profiled with CODEX (Supplementary Fig. 13, Pearson’s correlation coefficient between significance levels, *r* ≥ 0.98).

To identify molecular cues that potentially mediate the crosstalk between splenic cell populations, we compared differentially expressed genes to a database of receptor-ligand interactions^33^ and assessed the relative spatial location of ligand- and receptor-expressing cell populations. Overall, we detected 29 significant interactions based on this approach (Benjamini-Hochberg adjusted *q*-value ≤ 0.05, Supplementary Table 4), including the expression of several cues in red pulp macrophages related to the modulation of C1q-dependent phagocytosis, the F2L-mediated priming of neutrophils, and the IGF1-mediated activation of basophils (Fig. 4c, Supplementary Note). These interactions appeared substantially reduced or absent in monocyte-derived macrophages, suggesting the specialization of resident red-pulp macrophages in the positive regulation of humoral innate immune responses in the murine spleen. More broadly, the results of this analysis show the utility of spatially-resolved expression data in the study of extrinsic signaling between cell populations.

### Annotation of highly-multiplexed cytometry data using STvEA

Beyond the realm of mIHC, other technologies also allow for highly-multiplexed protein expression profiling of individual cells. Multi-parameter flow cytometry and cytometry by time-of-flight (CyTOF) provide high-dimensional proteomic characterizations of single-cell suspensions, and are frequently used in immunology and studies of cancer. We reasoned that the same procedure for annotating cell populations on mIHC images could be adapted to these modalities of data. To assess the utility of STvEA in this context, we first applied it to 114,568 wild-type mouse splenocytes profiled with CyTOF in a published study^2^. The panel in this study had 22 antibodies in common with our CITE-seq atlas. STvEA correctly consolidated the protein expression spaces of the CyTOF and CITE-seq datasets and annotated 13 cell populations (Fig. 5a), which represented 91% of the cells in the CyTOF dataset. As in our analysis of CODEX data, these annotations included subtle cell populations, such as pDCs and different stages of erythrocyte maturation, which are hard to identify without specifically tailored antibody panels. For comparison, we also performed more conventional analyses based on the algorithms X-shift, SPADE, and PhenoGraph, followed by manual annotation of the resulting clusters (Fig. 5b and Supplementary Fig. 9). Although the annotations produced by STvEA were consistent with the results of these analyses, STvEA provided an increase in the resolution and number of annotated cell populations with respect to manual annotations.

**Figure 5.**
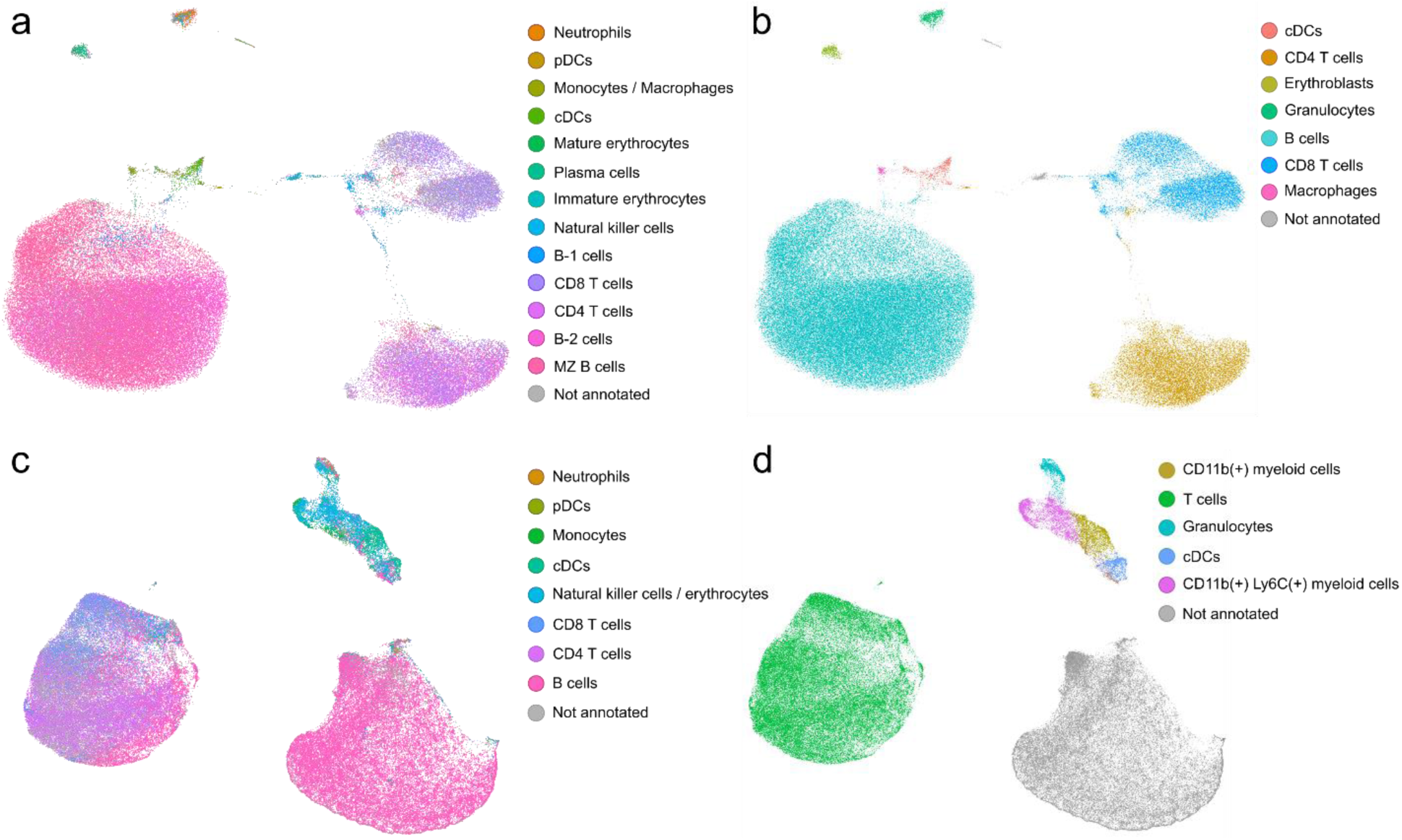
Transcriptome-guided annotation of mass cytometry data. **a)** UMAP representation of 114,568 mouse splenocytes profiled with CyTOF by Goltsev et al.^2^. The representation is labeled with the cell populations identified by STvEA based on a panel of 22 antibodies. **b)** The same representation is labeled according to the manually-annotated clusters produced by X-shift. Automated, transcriptome-guided annotations are consistent with manual analysis but provide an improvement in resolution and reproducibility. **c)** UMAP representation of 146,110 splenocytes from a glioma xenograft model profiled with CyTOF by Dusoswa et al.^34^. The representation is labelled with the cell populations identified by STvEA based on a panel of 11 antibodies. **d)** The same representation is labeled according to the manually-annotated clusters produced by X-shift. Annotation of this dataset is particularly challenging due to the small size of the antibody panel. However, although the annotations are limited in this case, STvEA still provides a substantial improvement with respect to standard procedures.

We next applied STvEA to 146,110 splenocytes from a glioma mouse model profiled with CyTOF^34^. We considered only 11 antibodies shared with our splenic CITE-seq atlas. There was a substantial overlap between the cell populations stained by these antibodies, making the annotation of this dataset particularly challenging. Despite these limitations, STvEA identified and annotated 8 phenotypically distinct cell populations (Fig. 5c), accounting for 82% of the cells in the CyTOF dataset. Although these annotations were broader than in other datasets we have analyzed, they still represented a substantial improvement with respect to procedures based on the manual annotation of clusters (Fig. 5d and Supplementary Fig. 9).

## Discussion

Methods for simultaneous profiling of protein and gene expression with single-cell resolution are evolving rapidly. Here, we have presented a computational approach for identifying and annotating cell populations in mIHC images by leveraging CITE-seq data of the same or closely related tissues. STvEA enables the optimal transfer of annotations from a CITE-seq dataset onto mIHC images or, more generally, highly-multiplexed cytometry data. We have demonstrated the utility of this approach with published mIHC and mass cytometry datasets of the murine spleen, and we have studied interactions between cell populations in this organ based on the inferred spatial patterns of gene expression.

Our work builds upon some of the recent developments in the integration of single-cell omics data^11, 12^ and is similar in spirit to previous studies mapping single-cell RNA-seq data to FISH images^35-39^. However, because mapping transcriptomic information onto cytometry data is carried out through the protein expression space, there are multiple conceptual and technical differences with those studies. Specifically, to consolidate protein expression measurements performed with multiple technologies (next-generation sequencing of oligo-tagged antibodies, imaging of fluorescently-labelled antibodies, and mass spectrometry of metal-tagged antibodies) we developed tailored normalization schemes for CODEX and CITE-seq. In addition, to account for the inaccuracy introduced by using relatively small antibody panels to map high-dimensional single-cell gene expression spaces, we proposed a new clustering approach to identify cell populations in the CITE-seq data that can be optimally mapped based on their protein expression profile.

STvEA is complementary to emerging technologies for simultaneous highly-multiplexed spatial profiling of proteins and transcripts, such as Digital Spatial Profiling^40^ (DSP) and Deterministic Barcoding in Tissue for Spatial Omics Sequencing^41^ (DBiT-seq). These technologies perform concurrent spatially-resolved proteomic and transcriptomic measurements in a tissue section and are therefore not subjected to mapping uncertainties. However, STvEA also provides a handful of unique advantages, such as its scalability to hundreds of thousands of cells (DSP can be only applied a few dozens of individual cells), its ability to work with sub-micrometer spatial resolutions (the spatial resolution of DBIT-seq is 20 μm), and its applicability to existing datasets, offering the possibility of performing new analyses of existing datasets. We therefore expect these tools to have complementary domains of applicability and be of great utility to researchers studying the cellular and molecular architecture of tissues, especially in light of the recent explosion of available single-cell RNA-seq and CITE-seq data of tissues.

## Methods

### Mouse handling

All animal work was approved by and carried out in compliance with the animal welfare regulations defined by the University of Pennsylvania International Animal Care and Use Committee (IACUC). 15 week old female BALB/cJ (Stock #000651) mice were acquired from The Jackson Laboratory (Bar Harbor, ME). Mice were allowed to age at the University of Pennsylvania Small Animal Facility until they reached approximately 9 months, at which point they were euthanized using CO2 followed by cervical dislocation.

### Tissue dissection and preparation of splenic single-cell suspensions

Spleens were removed from mice and mechanically dissociated with a syringe plunger over a 40 um strainer while being washed with 5 ml of PBS + 10% fetal calf serum. Suspensions were centrifuged briefly to pellet cells. Red blood cells were lysed with an RBC lysis buffer (155 mM NH4Cl, 12 mM NaHCO3, 0.1 mM EDTA) for 5 minutes and centrifuged again. 2 million cells from the resulting pellet were re-suspended in staining buffer (2% BSA, 0.01% Tween in PBS), and subsequently incubated with the antibody panel as described below (see “Cell Staining”).

### CITE-seq antibody conjugation and panel preparation

Antibodies were conjugated to 5’ amino-modified, HPLC-purified CITE-seq oligonucleiotides purchased from Integrated DNA Technologies. Antibodies were concentrated to 1 mg/ml in PBS pH 7.4 using 50 kDa cutoff spin columns (UFC505024, Millipore). Oligonucleiotides were resuspended to 1 mg/ml in 1x PBS pH 7.4 and were subsequently cleaned as suggested in the CITE-seq protocol. In brief, oligos were heated at 85°C and centrifuged at 17,000g to pellet any debris. For each antibody, 100 μg of antibody and 100 μg of oligo were conjugated using the Thunder-Link PLUS Oligo Conjugation System (SKU: 425-0300, Expedeon). All conjugates were cleaned as described in the CITE-seq protocol and resuspended to their final concentration in the Antibody Resuspension Buffer provided with the kit, with the exception of CD16/32, which was resuspended in 1x PBS. Successful conjugation was validated by running 1 μg of each conjugate on a 2% agarose gel which was subsequently stained with Sybr Gold (S11494, Thermo Fisher Scientific).

To prepare the panel, 1.5 μl of each antibody-oligo conjugate (except CD16/32) were combined in PBS and centrifuged in a 50 kDa cutoff column. After washing, the cleaned panel was recovered by flipping the column upside down and centrifuging. The cleaned panel was resuspended in staining buffer.

### Single-cell CITE-seq library preparation and sequencing

1.5 μg of the CD16/32 antibody-oligo conjugate was incubated with the single cell suspension for 10 minutes in place of the mouse seroblocker suggested in the CITE-seq protocol. The remaining 29 antibodies were then added to the cell suspension and incubated on ice. After incubation, cells were washed thoroughly, counted on a hemocytometer, and loaded into the 10x Chromium platform (10x Genomics) for single-cell library preparation. Cells were loaded at 1,200 cells/μl. Only samples with >80% cell viability were used, profiling a total of 2 mouse spleens. cDNA libraries were prepared following the standard CITE-seq and 10x protocols. The resulting antibody-derived tag (ADT) and mRNA libraries were combined at a 1:9 ratio and sequenced with an Illumina HiSeq 2500 at the Center of Applied Genomics, Children’s Hospital of Philadelphia.

### Multiplexed RNA FISH of splenic tissue sections

Whole spleens were removed from euthanized mice and immediately submerged in 4% paraformaldehyde for 5.5 hours. They were then cryoprotected in a 30% sucrose/70% fixative solution at 4°C until the tissue sank (approximately overnight, ∼16 hours). The tissue was embedded in OCT cryostat sectioning medium (OCT Compound, Sakura Finetek Inc, Supp. No. 4583) on dry ice and frozen at -80°C. Tissue was cut using a cryostat at -20°C into 10 μm-thick sections and frozen again at -80°C. Tissue was used for microscopy within 6 months of fixation and cryoprotection.

RNA fluorescence in situ hybridization experiments were carried out with the RNAscope Multiplex Fluorescence Reagent Kit v2 (Advanced Cell Diagnostics, Hayward, CA, USA, Cat. No. 323100). The RNAscope Assay for fixed frozen samples was followed per the manufacturer’s protocol with the following two modifications: the post-fix incubation was carried out with 4% PFA at room temperature for 90 minutes and manual target retrieval with a 5 minute sample incubation was performed instead of the steamer method. Probes for mouse *Bhlhe41* and *Il1b* (Advanced Cell Diagnostics, Cat. No. 467431 and 316891) were hybridized with Opal 520 (Akoya Biosciences, Cat. No. FP1487001KT), and probes for mouse *Cd79a* (Advanced Cell Diagnostics, 460181-C2) were hybridized with Opal 570 (Akoya Biosciences, Cat. No. FP1488001KT). Both dyes were diluted 1:1500 with TSA buffer provided by the RNAscope kit. Channel 2 was diluted in channel 1 1:50 as suggested in the RNAscope protocol. All incubations were carried out using a Stratagene PersonalHyb hybridization oven. Sequential sections were processed alongside the positive and negative controls provided by the RNAscope kit. Immunofluorescence images were acquired using a Leica TCS SP8 Multiphoton confocal microscope.

### Single-cell CITE-seq processing

We used Cell Ranger to de-multiplex, map to the mouse reference genome (mm10), and count UMIs in the mRNA libraries, and CITE-seq-Count to count UMIs in the ADT libraries. We filtered out cells with more than 10% UMIs from mitochondrially-encoded genes or less than 1,200 mRNA UMIs in total. We used scVI to infer a lower dimensional latent space for visualization and clustering of the mRNA expression data. scVI uses a neural network to fit a zero-inflated negative binomial model to represent the technical variation in scRNA-seq data and create a latent space. We inferred an 18-dimensional latent space representation for the expression data of all genes expressed in at least 15 cells (training size = 0.75, number of epochs = 400, learning rate = 1 × 10^−3^). The dimensionality of the latent space was empirically chosen based on the stability of the resulting representations and was consistent with the elbow of the scree plot. To visualize the mRNA expression, we further reduced the latent space to 2 dimensions using UMAP with Pearson’s correlation distance.

### Clustering and differential expression analysis of single-cell mRNA data

We clustered the cells in the latent space using HDBSCAN and an in-house consensus algorithm. Prior to clustering, we used UMAP to establish a metric in the 18-dimensional latent space, as suggested by the UMAP Python documentation. Then we scanned across the min_cluster_size and min_sample parameters of HDBSCAN (min_cluster_size ∈ {5,9,13,17}, min_sample ∈ {10,13,16,19,22,25,28,31,34,37}) and used cluster-based similarity partitioning to build a consensus matrix,

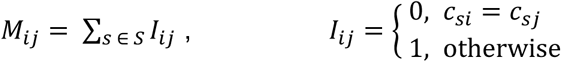

where *S* is the set of cluster assignments for all parameter configurations that gave rise to clusters with a silhouette score > 0.114, and *c*_*si*_ is the cluster ID of cell *i* in *s*. For this threshold, about 20% of the initializations of the UMAP metric and parameter scan do not produce any clusters with a satisfactory silhouette score. In this case, the UMAP and parameter scan were re-initialized with a new random seed. We used the above consensus matrix as a dissimilarity matrix among cells to produce a consensus clustering using average linkage agglomerative clustering (inconsistent value ≤ 0.1).

We ran edgeR’s general linear model (GLM) on the mRNA count data to identify differentially expressed genes between each cluster and all the other cells (fold change threshold > 2).

### Laplacian score analysis of single-cell mRNA data

We utilized the Laplacian score to more accurately annotate the mRNA data by identifying genes that have expression patterns within a cluster which cannot be explained by random variation. For each cluster, we built a graph where nodes represent cells and edges connect pairs of cells that are within ε distance, as defined by Pearson’s correlation in the latent space. We took ε to be given by the median pairwise distance among cells. For large clusters, we randomly sampled 1,000 cells. The Laplacian score *ℓ* of a gene with expression vector *f* is defined as

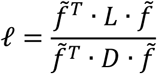

Where

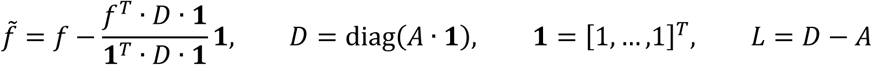

and *A* is the adjacency matrix of the graph. We computed the Laplacian score of the log(1 +*TPM* · 10^−2^) expression values for all genes expressed in at least 2% and at most 90% of the cells. To assess the significance of the Laplacian score as compared to random variation, we performed a permutation test by randomizing the cell labels 1,000 times.

### Normalization of ADT libraries

We fit the distribution of ADT counts for each antibody with a two-component negative binomial mixture model,

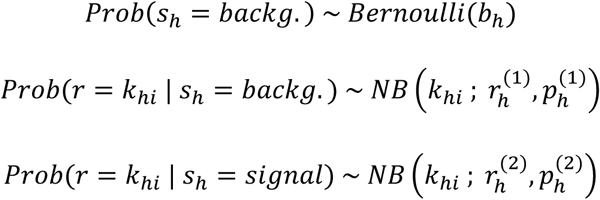

where *k*_*hi*_ represents the observed number of ADT UMIs for antigen *h* in cell *i*, the mixing parameter *b*_*h*_ represents the probability of a measurement of antigen *h* actually coming from the background, and the signal component is defined as the component of the mixture with highest median. Upon fitting the model using least-squares estimation, we filtered out the background component of the data by considering the matrix

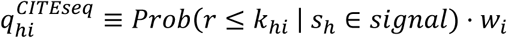

where the weights *w*_*i*_ are introduced to account for differences in the total number of ADT UMIs across cells,

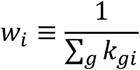

These weights can be justified by noting that *Prob*(*r*≤ *k*_*hi*_ | *s*_*h*_ ∈ *signal*) depends linearly on the total number of ADTs except at the tails of the distribution.

We performed batch correction on the 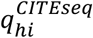 values using the mutual nearest neighbors approach of Haghverdi *et al.*^11^ and rescaled the resulting values to be in the [0,1] interval.

We did not consider CD169 in our analysis as it was showing expression in cell populations other than macrophages, possibly reflecting a lack of affinity of the conjugated antibody. In addition, the unimodal distribution of ERTR7 expression values was consistent with the fact that we did not capture stromal cells in our CITE-seq dataset (possibly because of the use of a non-enzymatic dissociation procedure). We therefore assigned all cells to the background component and set 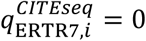.

### Processing of CODEX data

We considered the segmented and spillover-compensated CODEX data of the three wild-type mice profiled by Goltsev *et al.*^2^. We filtered out artifacts using a similar gating strategy to that of the CODEX protocol. We removed cells smaller than 1,000 or larger than 25,000 voxels. We then identified maximum and minimum cutoffs for blank channels by plotting the expression of one blank channel versus another, as described in the CODEX protocol. We removed cells with intensities above the upper cutoffs in any of the blank channels or below the lower cutoffs in all of the blank channels. Our cutoffs fell around the 99.5 and 0.2 percentiles respectively. However, we checked that small variations of the specific values did not greatly affect the number of cells removed.

### Normalization of CODEX data

We normalized the processed CODEX data by the total levels in each cell,

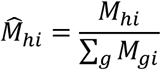

where *M*_*hi*_ is the level of antigen *h* in cell *i* before normalization. After this process, antigen levels are well approximated by a two-component Gaussian mixture model,

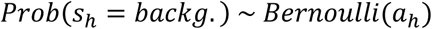

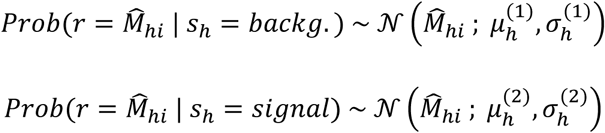

where the Gaussian with highest median corresponds to the signal component, and the mixing parameter *a*_*h*_ represents the probability of a measurement of antigen *h* actually coming from the background. Upon fitting the model to the data using the EM algorithm for maximum likelihood estimation, we filtered out the background component of the data by considering the probabilities

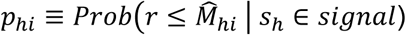

in subsequent analysis.

### Mapping of CODEX data into CITE-seq

We mapped the inferred CODEX probabilities *p* into the CITE-seq space *q*^*CITEseq*^ using a modified version of the general strategy proposed by Stuart *et al.*^*12*^. Specifically, we identified a set of anchors using a mutual nearest neighbors approach with *k*_anchor_ = 20. We found the nearest neighbors using Euclidean distance in a common 29-dimensional space obtained by canonical correlation analysis (CCA). We then filtered out anchors that do not preserve the structure of the original protein space. For that purpose, we kept only those for which the CODEX cell in the anchor was within the *k*_filter_ = 100 nearest CODEX cells to the CITE-seq cell in the anchor, or vice versa, as measured by Pearson’s correlation distance between *p* and *q*^*CITE*−*seq*^.

Cells in the CODEX dataset were aligned into the CITE-seq protein space using the following transformation

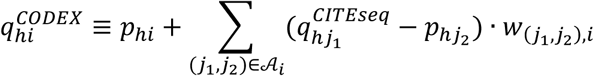

where 𝒜_*i*_ is the set of *k*_weight_ = 100 anchors (*j*_1_, *j*_2_) with smallest Pearson’s correlation distance between vectors 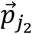 and 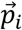 (with components 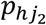 and *p*_*hi*_, respectively), and 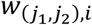 are weights specifying the effect size of anchor (*j*_1_, *j*_2_) on the CODEX cell *i* based on both mRNA and protein data,

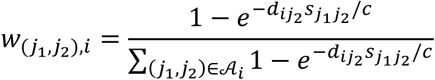

In this equation, 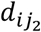 denotes Pearson’s correlation distance between the vectors 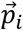 and 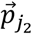, and *c* is a parameter specifying the width of the Gaussian kernel. The number of shared neighbors between the two anchor cells, 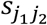, is defined as

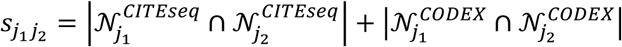

where 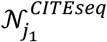 is the set of nearest CITE-seq cells to cell *j*_1_ in the mRNA latent space, 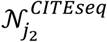 is the set of nearest CITE-seq cells to cell *j*_2_ in the CCA space, 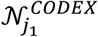 is the set of nearest CODEX cells to cell *j*_1_ in the CCA space, and 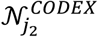 is the set of nearest CODEX cells to cell *j*_2_ in the CCA space. As before, distances in the mRNA and CCA spaces were measured using Pearson’s correlation and Euclidean distance respectively. In all cases, the number of nearest neighbors was chosen to be *k*_score_ = 80. The values 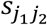 were scaled such that the 0.9 quantile is at 1 and the 0.01 quantile is at 0, and values above or below these quantiles were set to 1 or 0 respectively.

Since randomly sampled sections of the CODEX dataset can be mapped independently and concatenated later, we divided the CODEX dataset into 8 random sections of the same size (9,900 cells) to provide a speed improvement for the nearest neighbor calculations.

To be able to transfer quantities between the CITE-seq and CODEX datasets, we then built a ℳ^*CITEseq*→*CODEX*^ transfer matrix,

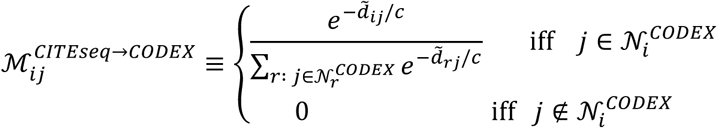

where 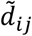 denotes Pearson’s correlation distance between the vectors 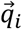 and 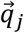 (with components 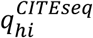 and 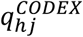, respectively), and *c* is a parameter that specifies the width of the Gaussian kernel. The set 𝒩_*i*_^*CODEX*^ contains the nearest CODEX cells to the CITE-seq cell *i* as measured by 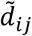, where *k*_transfer_ ≡ |𝒩_*i*_^*CODEX*^| = 0.002 × *n*_*CODEX*_, and *n*_*CODEX*_ is the number of cells in the CODEX dataset. These matrices can be used to transfer quantities across the two datasets. For instance, the inferred mRNA expression level of gene *m* in the CODEX cell *j* is given by

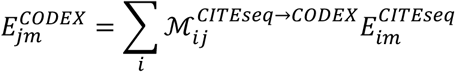

where *E*^*CITEseq*^ denotes the mRNA expression matrix in the CITE-seq dataset. Similarly, the mRNA cell populations can be mapped to the CODEX data using

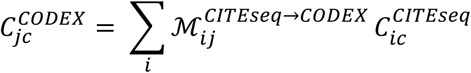

where the sum runs over all cells in the CITE-seq dataset and 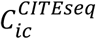 is the indicator function of cluster *c*. Note that due to the mapping uncertainties, the resulting feature vector is no longer a binary vector. To assess mapping uncertainties (Fig. 2c), we computed the Pearson’s correlation coefficient of the vectors 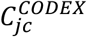 that result from restricting the above sum to cells in each of the two mice profiled with CITE-seq.

### Parameter selection

For different values of *k*_anchor_, *k*_filter_, and *k*_score_, we evaluated the performance of the algorithm to accurately map a set of “gold standard” cell populations. The populations we considered were B cells, T cells, NK cells, dendritic cells, neutrophils, plasma cells, and red-pulp macrophages, as they were general enough to be clearly identifiable in both datasets by clustering and the expression of specific markers. We used the Louvain community detection algorithm in a *k* = 49 nearest neighbor graph for clustering the CODEX protein data. To quantify the performance of the mapping, we defined the quality scores of a set of anchors 𝒜 as

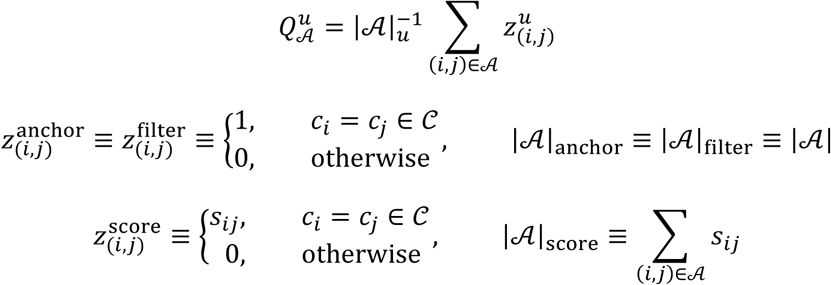

where *c*_*i*_ is the cell type of cell *i* and 𝒞 is the set of “gold standard” populations. We sequentially chose the values of *k*_anchor_, *k*_filter_, and *k*_score_ that maximized these quality scores.

### Quantification of mapping uncertainties and stability

To study the consistency between CODEX cells that are mapped by the same CITE-seq cell, we randomly selected *k*_transfer_ pairs of cells among the CODEX cells that map to the same CITE-seq cell, and calculated Person’s correlation between antigen levels in each pair of cells. The mean value of the correlation coefficients was taken to represent the mapping uncertainty of each CITE-seq cell, which was shown on the UMAP in the Supplementary Fig. 5. To compare these correlation coefficients with the correlation between two random pairs of CODEX cells, we considered 7,097 random sets of *k*_transfer_ CODEX cells and computed the correlation coefficients using the same method. To assess localized uncertainty in the mapping algorithm, we defined a mapping score for each CODEX cell as 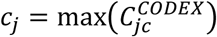 and plotted these scores on the UMAP representation of the CODEX protein space (Supplementary Fig. 6).

To study how the size of the CITE-seq dataset affects the performance of STvEA, we randomly sampled 5000, 2500, and 1000 cells from the original CITE-seq dataset, and ran STvEA using the same default parameters.

To assess the effect of antibody panel selection on STvEA mapping, we used the glmnet R package^42^ to perform multinomial logistic lasso regression on the CITE-seq mRNA clusters with respect to the protein expression levels. We identified values of the regularization parameter λ for which a subset of *n* = 25, 21, …, 13, 9 antibodies had non-zero coefficient and ran STvEA truncating the antibody panel to each of these subsets. The stability of STvEA was evaluated for each antibody panel by computing the Pearson’s correlation coefficient between the vectors of cell population assignments, 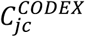, for each CODEX cell when using the full and a reduced antibody panel (Supplementary Fig. 7).

### Optimized annotation of cell populations

To annotate mIHC images with STvEA while accounting for uncertainty in the mapping algorithm, we sought to identify a clustering of the CITE-seq cells that allows for the highest resolution in number of clusters while maintaining a good modularity of the clusters upon mapping into the CODEX protein expression space. We started from the simplified hierarchical tree of an HDSBCAN clustering that passed the silhouette threshold in the original CITE-seq consensus clustering (*min_cluster_size* = 17, *min_samples* = 34). For computational simplicity, each CODEX cell was assigned to its closest CITE-seq neighbor in the shared protein space *q*. Each branch of the hierarchical tree could then be easily mapped onto CODEX. Since HDBSCAN allows some cells to remain unclustered, each bifurcation in the tree may have some “singlets” that are contained in the parent cluster but not the child clusters. To fill out the tree, we imputed these singlets into the closest child cluster based on Pearson’s correlation distance in the protein space. We then implemented an agglomerative clustering approach based on this tree by computing the modularity in the CODEX protein expression space. We considered the *k* = 50 nearest neighbor graph generated by Pearson’s correlation distance between the protein expression profiles of the CODEX cells. For each bifurcation in the tree, we computed the subgraph spanned by the cells in the two clusters involved in the bifurcation, and then computed the Newman-Girvan modularity^43^ (also known as Louvain modularity) of the clusters in this subgraph. Starting at the leaves of the tree, two child clusters were merged into their parent if the modularity of the bifurcation was less than a quality threshold *t*_*q*_, or if the modularities between the cells in each child cluster versus all other cells in the CODEX dataset were less than a sparsity threshold *t*_*s*_ (*t*_*q*_ = 0.1, *t*_*s*_ = 0.003). In cases where a single-cluster branch in a bifurcation did not pass the sparsity threshold, the cells from this cluster where merged into sibling clusters. Similarly, cells of a single-cluster branch were merged into sibling clusters if this represented an increase in modularity larger than an imputation threshold *t*_*i*_ (*t*_*i*_ = 5%). After this process, we smoothened the cell assignments in CODEX based on each cell’s neighbors. For each CODEX cell *x*, we defined a neighborhood of other cells *N*_*x*_ within a protein expression correlation threshold (Pearson > 0.99). The cell assignment of a cell *x* is then defined as,

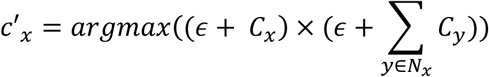

where *C*_*x*_ is the indicator function of the cluster of *x*, and *ϵ* is a constant to control the contribution of neighboring cells (*ϵ* = 0.01).

### Spatial relationship among cell populations

To assess the spatial relationship between two feature vectors *f* and *g* defined over the cells in the CODEX dataset, we built a *k* = 2 nearest neighbor graph using Euclidean distance in the CODEX spatial dimensions expressed in nm (median distance 5 μm). We then introduced the adjacency score, defined as

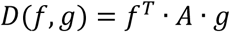

where *A* is the adjacency matrix of the nearest neighbor graph. This score takes high values when the features take high values in adjacent cells. The scale of the interactions is set by the magnitude of the nearest neighbor parameter *k*. Features that we have used in this paper include cell population assignments 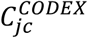 (to assess whether two cell populations co-localize spatially) and mapped gene expression 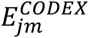 (to assess whether genes encoding for ligands and receptors are expressed in adjacent cells). The significance of this score was assessed using a null distribution built by permuting the cell ID’s. For mutually exclusive binary features (such as cluster assignments) the null distribution can be computed analytically in terms of the hypergeometric distribution Hypergeom(*u*; *N, K, n*),

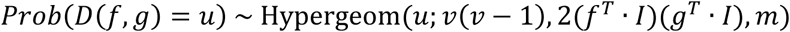

where *v* and *m* are the number of nodes and edges in the nearest neighbor graph, respectively, and *I* is the identity matrix. For non-binary features, we did not find a closed form for the null distribution, so we approximated it using a normal distribution whose parameters were estimated from 1,000 random permutations. We controlled the false discovery rate for multiple hypothesis testing using the Benjamini-Hochberg *q*-value procedure.

To account for the effect of mapping uncertainties on the adjacency score of cell populations, we also computed the overlap score, *f*^*T*^ · *g*, and assessed its significance by randomly permuting the entries of one of the feature vectors. In addition, we evaluated the Pearson’s correlation of the adjacency score *q*-values across the three mice profiled with CODEX (Fig. 3b).

### Identification of paracrine interactions

We used CellPhoneDB^33^ to identify significant ligand-receptor pairs within the CITE-seq mRNA expression data. CellPhoneDB identifies genes coding for ligand and receptor pairs that are differentially expressed in one or more cell populations using a curated database of ligands and receptors. Since CellPhoneDB only considers human gene pairs, we generated a mouse ortholog database of ligands and receptors using Ensembl^44^ (version 96). For simplicity, this analysis was restricted to only those genes which have a unique ortholog. The 67 significant interactions (CellPhoneDB *p*-value ≤ 0.05) identified by this analysis were then filtered using the adjacency score approach described above (see “Spatial relationship among cell populations”) to identify pairs of genes significantly expressed in adjacent cells. The expression of any complexes output by CellPhoneDB was calculated as the sum of the expression of their component genes.

### Analysis of CyTOF data

We applied STvEA to the CyTOF datasets of Goltsev et al.^2^ and Dusoswa et al.^34^. The dataset of Goltsev et al. consists of 124,277 cells in total and has 22 antibodies in common with our CITE-seq panel. To preprocess this data, we first removed outlier cells that express fewer than 500 total counts in the 22 antibodies or are within the top 2% of cells for total counts. We applied an arcsinh transform (cofactor 5) to the remaining 114,568 cells and scaled the resulting values to the interval [0,1]. We use STvEA as described above to map our CITE-seq atlas into this dataset. We then applied the optimized clustering approach described above (*t*_*q*_ = 0.2, *t*_*s*_ = 0.001, *t*_*i*_ = 3.5%) to identify 16 clusters in the CyTOF dataset associated with 13 phenotypically different cell populations in the CITE-seq atlas.

For the dataset of Dusoswa et al., we considered all 146,119 cells passing the “Live singlets” gate from the MGL02_Spleen sample. We removed cells with zero expression in the 11 antibodies considered in our analysis, resulting in 146,110 cells in total. We scaled the arcsinh transformed (cofactor 5) values to the interval [0,1], and applied STvEA as described above to map our CITE-seq atlas onto this dataset. We used the optimized clustering approach described above (*t*_*q*_ = 0.25, *t*_*s*_ = 0.001, *t*_*i*_ = 1%) to identify 23 clusters in the CyTOF dataset associated with 8 phenotypically different cell populations in the CITE-seq atlas.

### Annotation of X-shift, SPADE, and PhenoGraph clusters using StvEA

We used the implementations of these algorithms in the VorteX Java software (https://github.com/nolanlab/vortex), the PhenoGraph Python package (https://github.com/jacoblevine/PhenoGraph), and the SPADE R package (https://github.com/nolanlab/spade). For all datasets, we ran the algorithms using default parameters and restricted to those cells used in our STvEA analysis. To create the pie chart graphs in Supplementary Fig. 9, we started with the output graph produced by SPADE, or we created one from the X-shift and PhenoGraph clusters by computing a minimum spanning tree between cluster centroids using the Pearson’s correlation distance between the protein expression profiles. For each node in the graph, we identified the proportion of cells from each mapped CITE-seq cluster (see “Optimized annotation of cell populations” above) in that node, and visualized those proportions as a pie chart.

### Online database

The complete results of our analysis can be interactively queried through a web application hosted at the URL: https://camara-lab.shinyapps.io/stvea.

### STvEA software

All algorithms have been implemented and documented in an R package. The package can be downloaded from the URL: https://github.com/CamaraLab/STvEA.

## Acknowledgements

The authors would like to thank Daniel Aldea, Rachael Aubin, Jake Crawford, Doug Epstein, Yugong Ho, Jesse Humenik, Yana Kamberov, Steve Liebhaber, Fernanda Mafra, Michael Peel, Renata Pellegrino, Qi Qiu, Staci Rakowiecki, Jacqueline Smiler, Ryan Staupe, John Wherry, Hao Wu, and the Zhou lab for scientific discussions and assistance with various aspects of the experiments. The work of P.G.C. and S.W. is partially funded by Stand-Up-To-Cancer Convergence 2.0.

## Author Contributions

E.C.T. performed all the experiments. K.W.G. implemented all the algorithms and performed the analyses. Z.M. assisted with the analyses. S.W. designed and implemented the web application. P.G.C. conceived the study and supervised the work. All authors contributed to writing the manuscript.

