## Supplementary Material for "Single-Cell Transcriptomic Analysis of mIHC Images via Antigen Mapping"

### Supplementary Note

The results of our analysis suggest a specialization of resident red pulp macrophages in regulating the homeostasis of humoral innate immune responses in the spleen (Fig. 4c). We identified the expression of several cues related to the positive regulation of these responses. Complement component 1q (C1q) can mediate phagocytosis of apoptotic cells by calreticulin/LRP1 receptor stimulation[1]. We observed the co-localized expression of genes encoding for C1q subunits and the receptor LRP1 in both red pulp and monocyte-derived macrophages (Fig 4c and Supplementary Table 4). However, the expression of C1q genes was 12-fold higher in red pulp macrophages than in monocyte-derived macrophages ($p$-value < $10^{-10}$), suggesting the specialization of the former in modulating C1q-dependent phagocytosis. Red pulp macrophages also displayed high expression levels of the *Hebp1* gene, which encodes for the precursor of the F2L peptide that activates and chemo-attracts neutrophils by binding to the formyl peptide receptor Fpr2[2]. Consistent with a function of red pulp macrophages in orchestrating neutrophil activation, *Hebp1*-expressing red pulp macrophages were in close spatial proximity to *Fpr2*-expressing neutrophils (Fig. 4c, Supplementary Table 4). In contrast, monocyte-derived macrophages displayed low expression levels of *Hebp1* (fold-change with respect to red pulp macrophages = 0.1, $p$-value < $10^{-10}$). These macrophages instead expressed annexin A1 (*Anxa1*) (Fig. 4c, Supplementary Table 4), an anti-inflammatory agonist of the neutrophilic receptor Fpr1[3]. Additionally, we observed high expression levels of insulin-like growth factor 1 (*Igf1*) in red pulp macrophages. Various works have put forward the role of human IGF-1 in stimulating the activation and chemokinesis of basophils through the IGF-1R receptor[4-7]. Consistent with this hypothesis, our analysis revealed red pulp macrophages were recurrently adjacent to *Igf1r*-expressing basophils (Fig. 4c, Supplementary Table 4). The expression of *Igf1* in monocyte-derived macrophages was substantially lower (fold-change with respect to red pulp macrophages = 0.1, $p$-value < $10^{-10}$), indicating the specialization of resident red pulp macrophages in IGF-1 signaling. Hence, taken together these results suggest important differences in the immuno-regulatory function of red pulp and monocyte-derived macrophages in the murine spleen.

**Supplementary Figures**

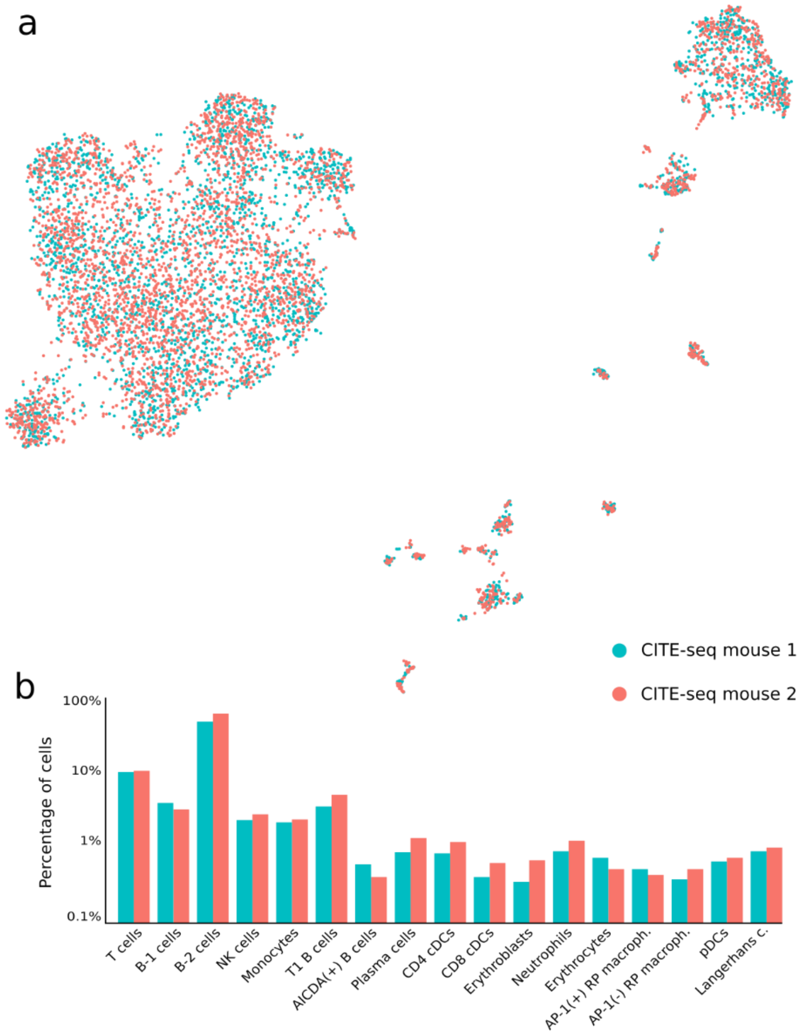


**Supplementary Figure 1. Absence of substantial batch effects in the CITE-seq atlas of the murine spleen. a)** The UMAP representation of the mRNA expression data is labelled according to the two murine spleens that were profiled with CITE-seq. **b)** Percentages of the cells of each profiled mice assigned to each cluster.

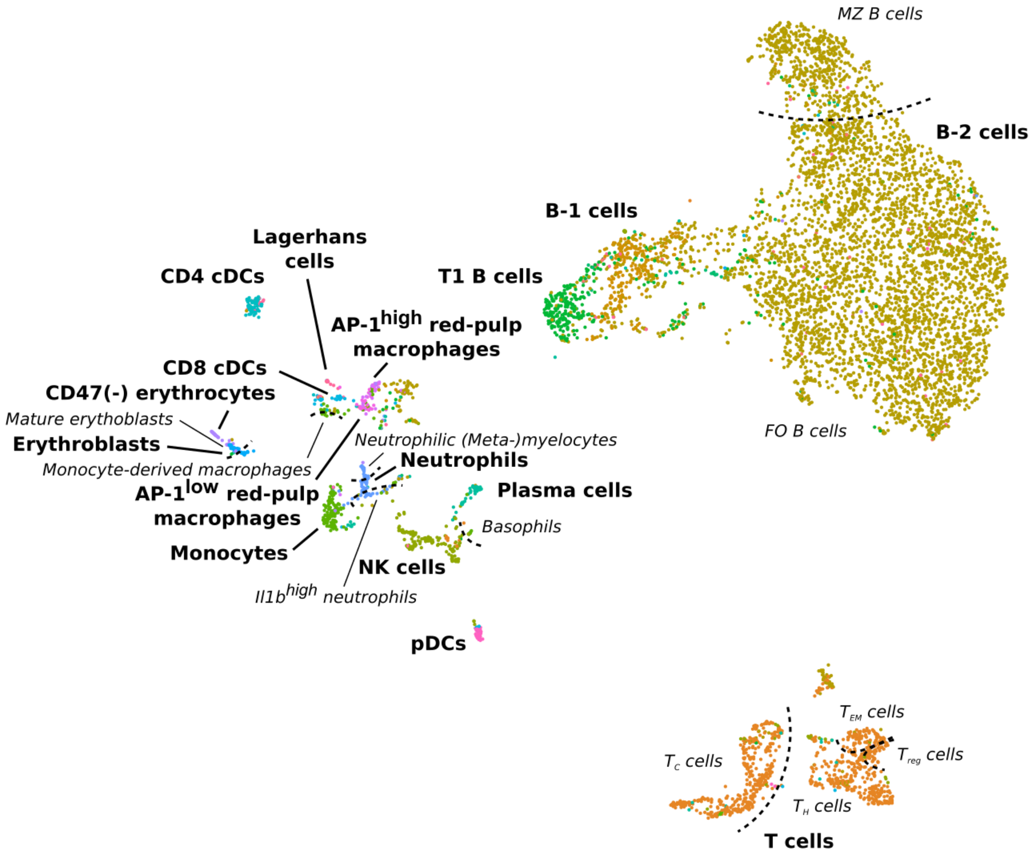


**Supplementary Figure 2. UMAP representation of the protein expression data of 7,097 cells from the murine spleen profiled with CITE-seq.** Cells are labelled according to the cell populations identified from the mRNA expression data.

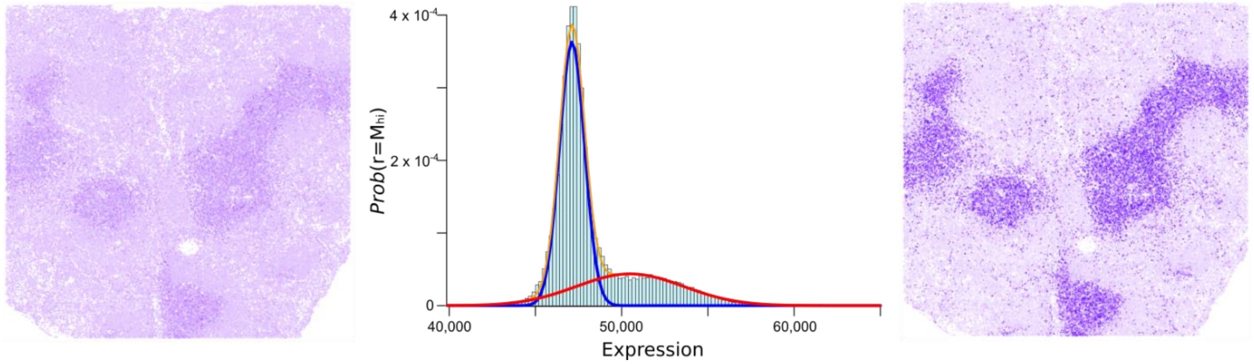


**Supplementary Figure 3. Normalization of CODEX data using a Gaussian mixture model.** The levels of CD4 protein determined by CODEX are shown in a splenic section after standard processing of the data (left). A two-component Gaussian mixture model was fit to the CD4 protein levels, where the lowest and highest mode components correspond to background and signal, respectively (middle). Upon filtering out the background component, the CD4 signal at T-cell follicles becomes more evident.

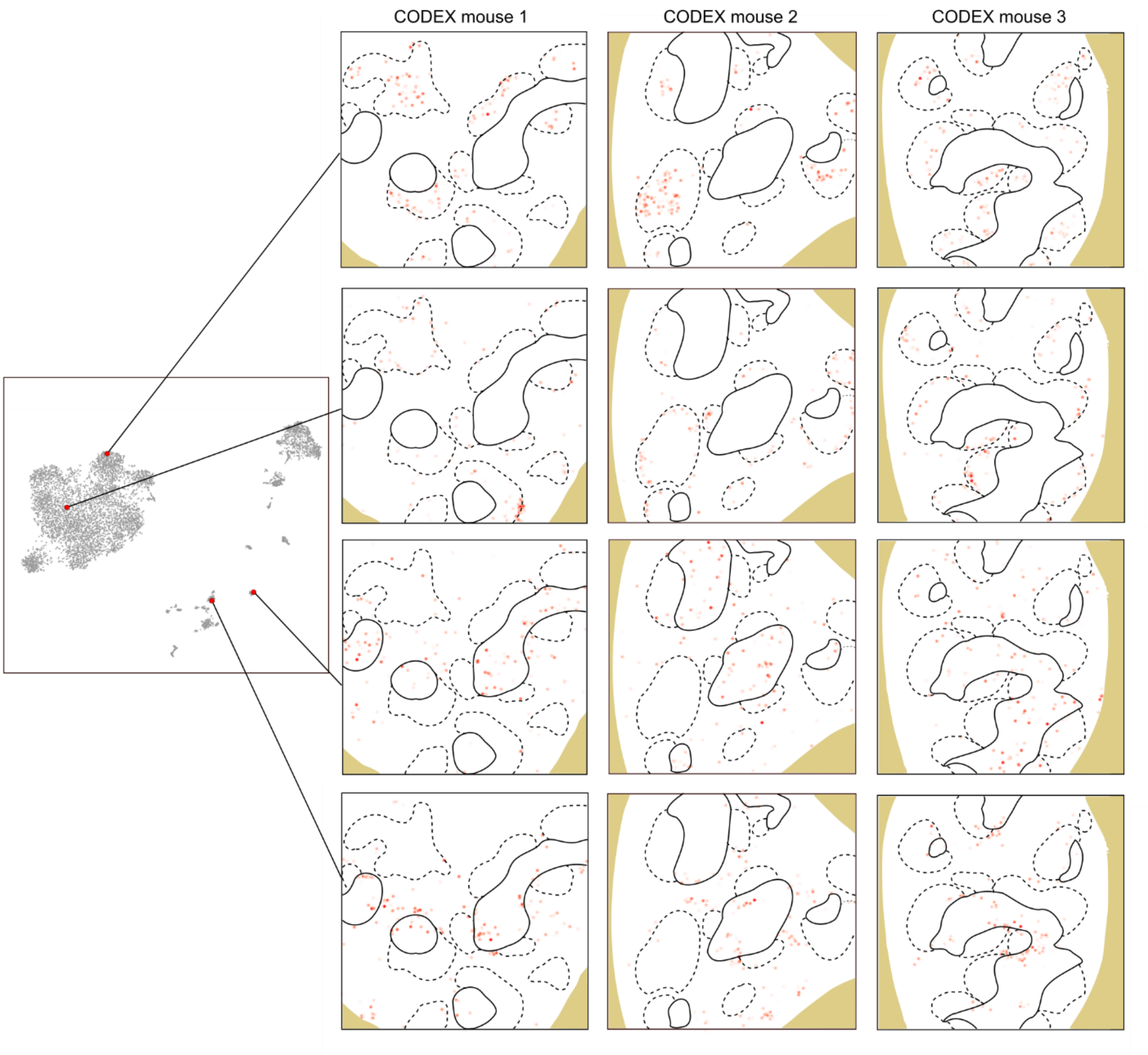


**Supplementary Figure 4. Consistency of the inferred locations of cells from the splenic CITE-seq atlas across 3 different splenic sections profiled by CODEX.** The inferred spatial locations of 4 different cells (two B-2 cells, one pDC, and one CD4 cDC) are shown in the 3 splenic sections, showing a high degree of consistency in their relative location with respect to the T and B cell zones (indicated by continuous and dashed lines, respectively).

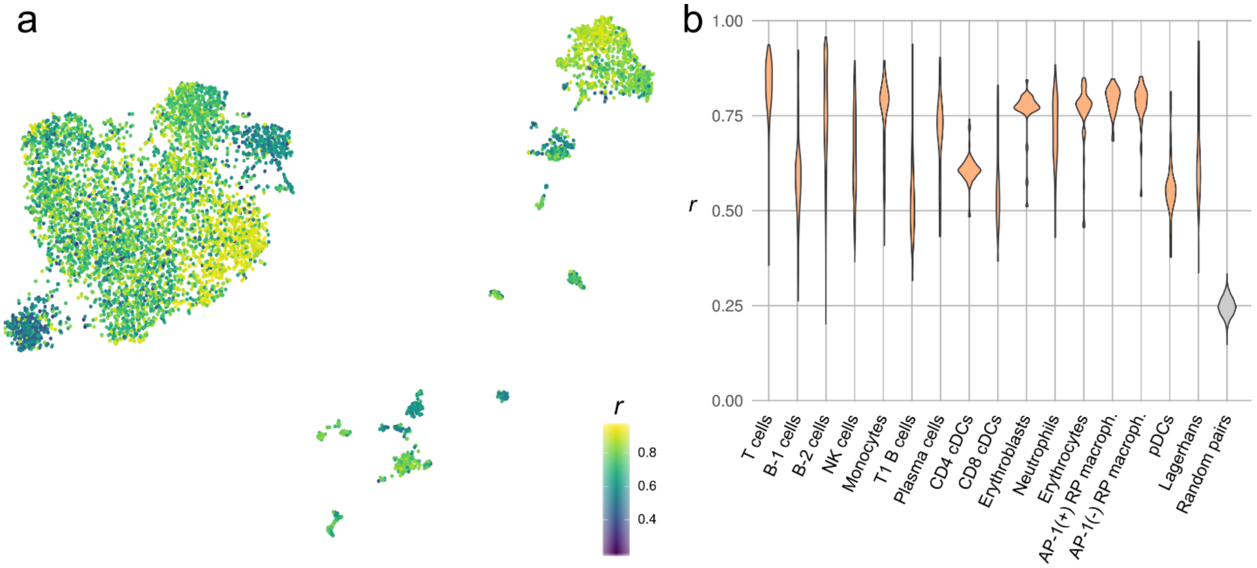


**Supplementary Figure 5. Consistency of protein expression profiles of CODEX cells related to the same CITE-seq cell. a)** Each CITE-seq cell in the UMAP representation of the mRNA expression data is labeled by the average Pearson's correlation coefficient between the protein expression profiles of all CODEX cells related to that cell. The level of correlation varies across the CITE-seq dataset, reflecting differences in mapping uncertainty. **b**) Violin plot of the average Pearson's correlation coefficient between the protein expression profiles of CODEX cells related to the same CITE-seq cell, grouped by cell type. CODEX cells that map to B-1 B cells, T1 B cells, and dendritic cells have the lowest correlation coefficients. However, for all cell types correlation coefficients are substantially larger than those between the protein expression profiles of random pairs of CODEX cells (represented in grey).

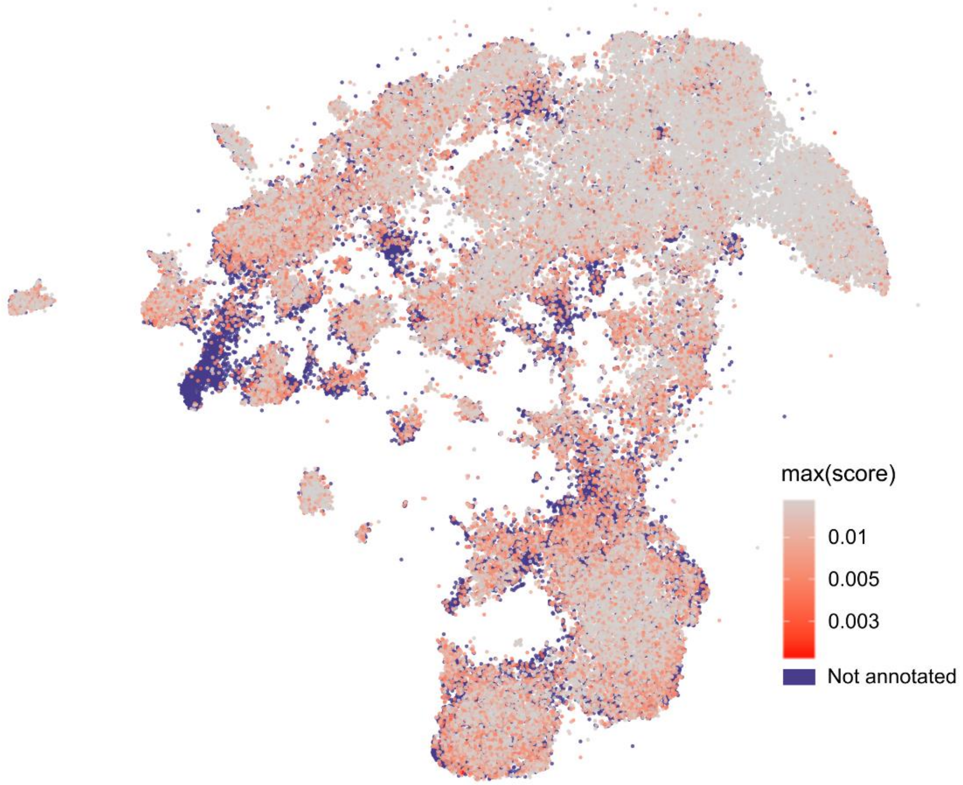


**Supplementary Figure 6. Mapping scores across the CODEX dataset.** The UMAP representation of the CODEX protein expression space is labeled by the maximum mapping score, $c_j = \max\left(C_{jc}^{CODEX}\right)$ (see Online Methods). Cells with low mapping scores are more sensitive to the size of the CITE-seq dataset.

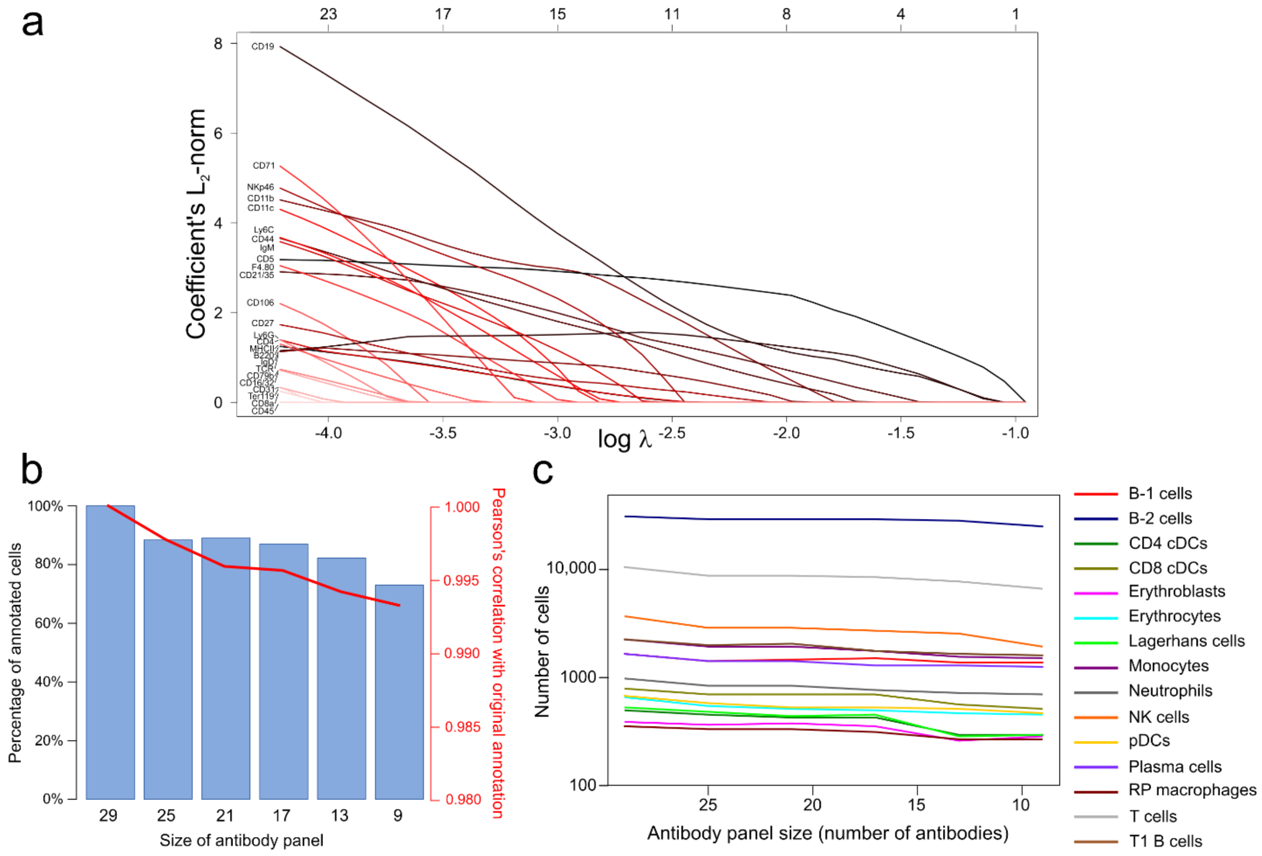


**Supplementary Figure 7. Dependence of STvEA on the size of the antibody panel.**
**a)** Multinomial logistic lasso regression on the CITE-seq mRNA clusters of the splenic atlas with respect to the protein expression levels. The $L_2$ norm of the regression coefficient of each antibody is represented as a function of the regularization coefficient. This analysis allows us to rank the antibodies according to their predictive value and build smaller-size antibody panels. **b)** Percentage of annotated CODEX cells in the original STvEA analysis of the murine spleen that are still annotated when using a smaller antibody panel. The median Pearson's correlation coefficient between the annotations in the analyses using the full and reduced antibody panel is also shown. **c)** Number of CODEX cells for each annotated cell population as a function of the antibody panel.

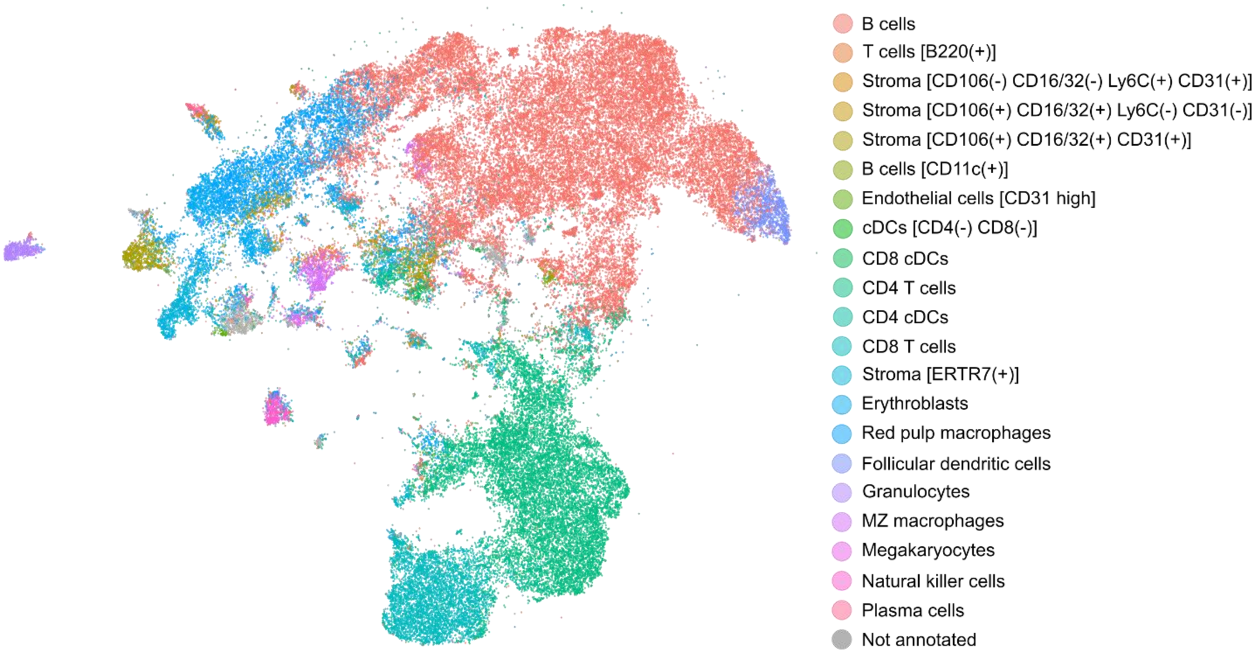


**Supplementary Figure 8. Manual annotation of the murine spleen CODEX dataset.** The UMAP representation of the CODEX protein expression space is labeled according to the manually annotated X-shift clusters of Goltsev et al.

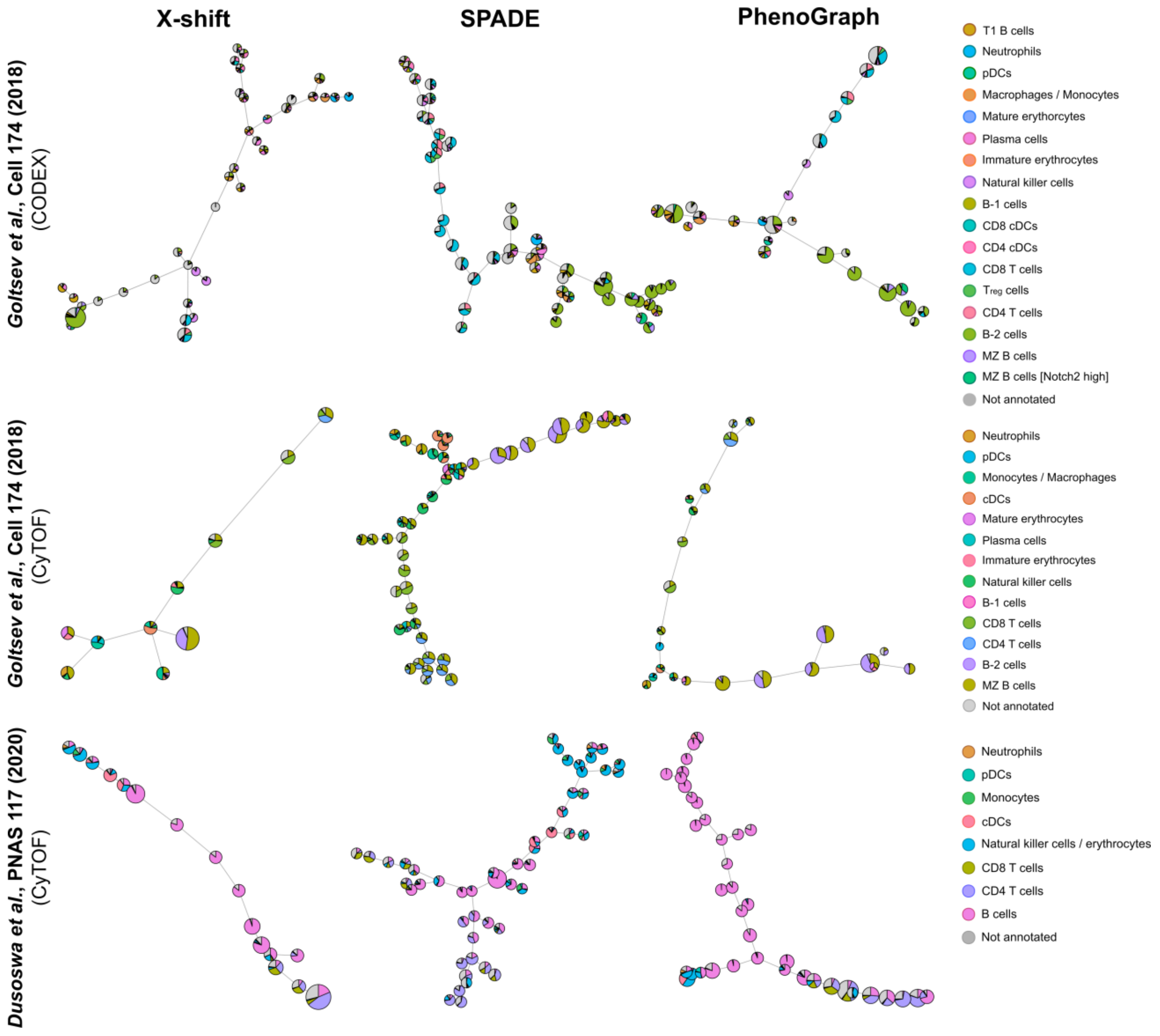


**Supplementary Figure 9. STvEA-based annotation of clusters produced by cytometry data clustering algorithms.** Minimal spanning tree representations of the clustering produced by X-shift, SPADE, and PhenoGraph on published CODEX and CyTOF datasets of the murine spleen. Each cluster is represented by a pie chart depicting its composition in terms of STvEA cell population assignments.

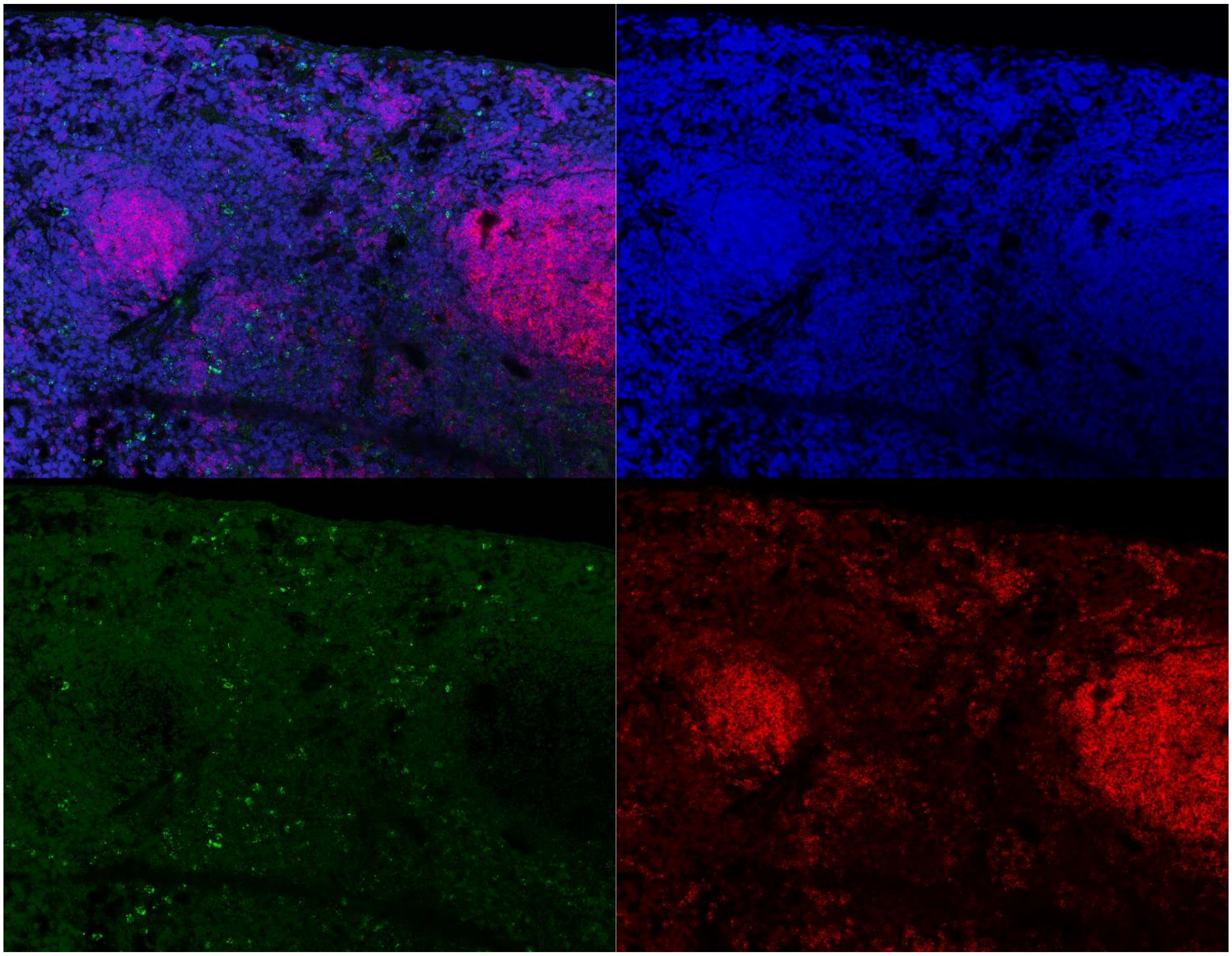

**Supplementary Figure 10. RNA FISH of the murine spleen depicting DAPI (blue), *Il1b* (green), and *Cd79a* (red).**

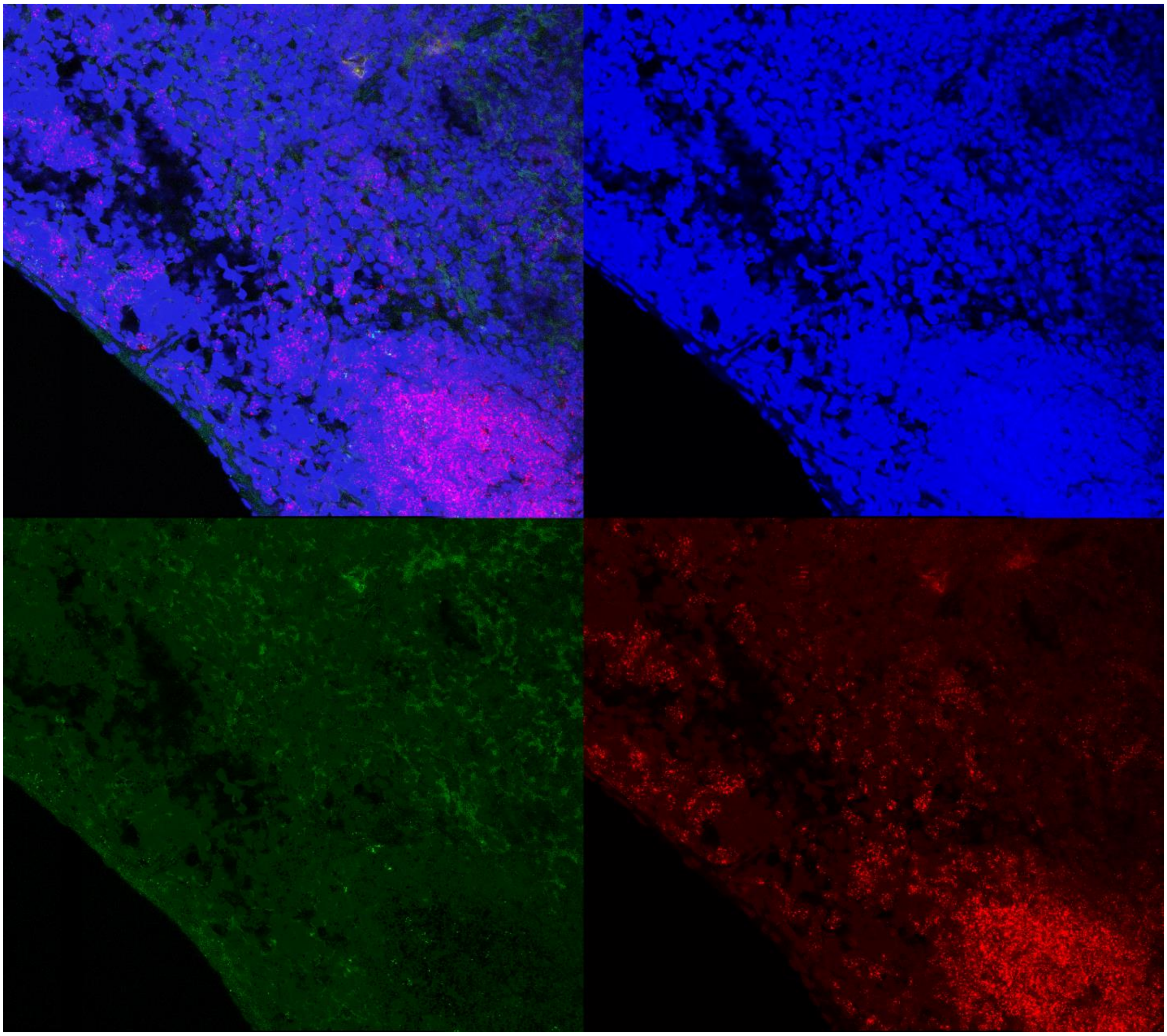

**Supplementary Figure 11. RNA FISH of the murine spleen depicting DAPI (blue), *Bhlhe41* (green), and *Cd79a* (red).**

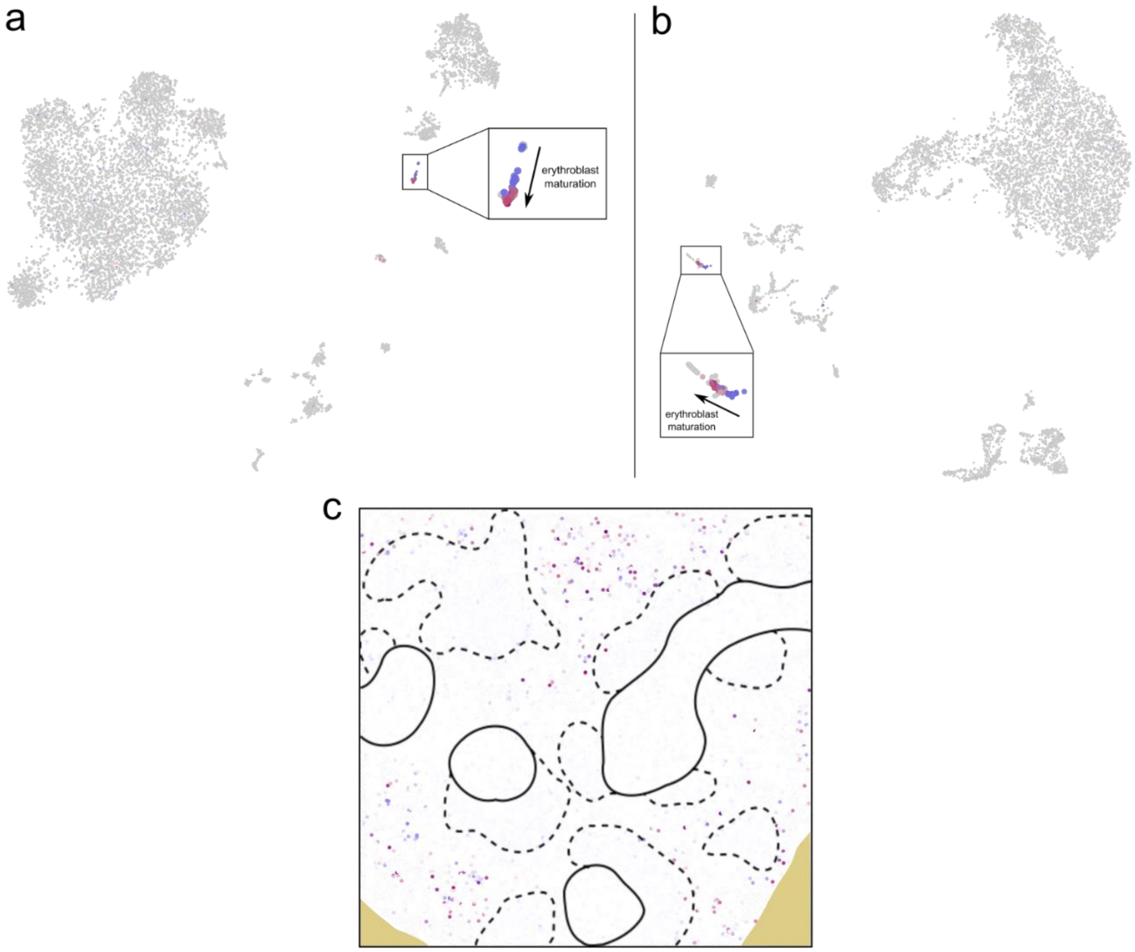


**Supplementary Figure 12. Analysis of splenic erythroblast maturation with STvEA.** The mRNA expression levels of *Car1* (blue), a marker of early stages of erythropoiesis, and *Gypa* (red), a marker of intermediate and late stages of erythropoiesis, are depicted in the UMAP representations of the mRNA (a) and protein (b) CITE-seq data, and mapped to a splenic tissue section profiled with CODEX (c).

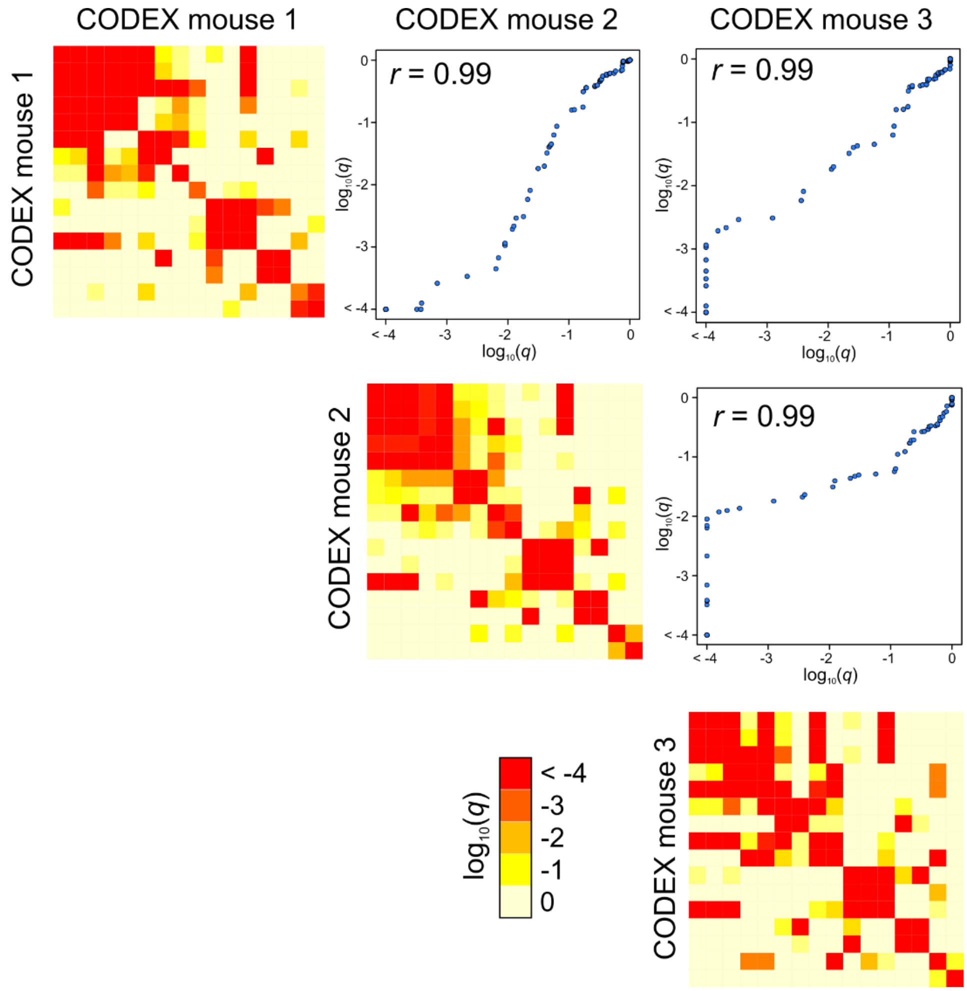


**Supplementary Figure 13. Consistency of the analysis of cell-to-cell interactions across different splenic sections profiled by CODEX.** Three different splenic sections were independently mapped to the CITE-seq atlas. The inferred spatial co-localization patterns are highly consistent across the three sections, as indicated by the high Pearson's correlation coefficients among the estimated significances.

### Supplementary Tables

| Target | Clone | Catalog Number | Manufacturer |
|---|---|---|---|
| CD45/B220 | RA3-6B2 | 553084 | BD |
| CD106 | 429(MVCAM.A) | 553330 | BD |
| CD11b | M1/70 | 553308 | BD |
| CD11c | H13 | 117302 | Biolegend |
| CD16/32 | 2.4/G2 | 553142 | BD |
| CD169 | MOMA-1 (MCA947) | MCA947G | Bio-rad |
| CD19 | 1D3 | 553783 | BD |
| CD21/35 | 7G6 | 553817 | BD |
| CD27 | LG.3A.10 | 124202 | Biolegend |
| CD3 | 17A2 | 555273 | BD |
| CD31 | MEC13.3 | 553370 | BD |
| CD35 | 8C12 | 558768 | BD |
| CD4 | RM4-5 | 553043 | BD |
| CD44 | IM7 | 553131 | BD |
| CD45 | 30-F11 | 553076 | BD |
| CD5 | 53-7.3 | 100602 | Biolegend |
| CD71 | C2 | 553264 | BD |
| CD79b | HM79 | 132802 | Biolegend |
| CD8a | 53-6.7 | 553027, BD | BD |
| CD90 | G7 | 105202 | Biolegend |
| ER-TR7 | | MCA2402 | Bio-rad |
| F4/80 | T45-2342 | 565409, BD | BD |
| IgD | 11-26c.2a | 553438, BD | BD |
| IgM | R6-60.2 | 553405, BD | BD |
| Ly-6C | HK1.4 | 128001 | Biolegend |
| Ly-6G | 1A8 | 551459 | BD |
| MHCII I-A/I-E | M5.114.15.2 | 556999 | BD |
| Ter-119 | TER-119 | 116202 | Biolegend |
| NKp46 | | 560754 | BD |
| TCRbeta | H57-597 | 553167 | BD |

**Supplementary Table 1. List of antibodies used for the single-cell CITE-seq atlas of the murine spleen.** We used the same clones and vendors in our panel as in Goltsev et al.[8].

**Supplementary Table 2. Differentially expressed genes in the single-cell transcriptomic analysis of the murine spleen.** For each cell population, the differentially expressed genes with respect to the combined other cell populations are presented. [Provided as a separate file].

**Supplementary Table 3. Laplacian score analysis of the splenic single-cell transcriptomic data.** For each cell population, the Laplacian score for each gene is presented. Genes with small scores have significant transcriptional heterogeneity within that population. [Provided as a separate file].

**Supplementary Table 4. Analysis of ligand-receptor gene interactions.** Candidate interactions based on the expression of genes coding for ligands and receptors. For each gene, the spatial proximity of the ligand and receptor expression was assessed through the adjacency score. [Provided as a separate file].
